# Schrödinger’s range-shifting cat: analytic predictions for the effect of asymmetric environmental performance on climate change responses

**DOI:** 10.1101/2022.02.04.479140

**Authors:** J. Christopher D. Terry, Jacob D. O’Sullivan, A. G. Rossberg

## Abstract

Analytic models for how species will respond to climate change can highlight key parameter dependencies. By mapping equations for population dynamics onto corresponding well-studied problems from quantum mechanics we derive analytical results for the frequently observed case of asymmetric environmental response curves. We derive expressions in terms of parameters representing climate velocity, dispersal rate, maximum growth rate, niche width, high-frequency climate variability and environmental performance curve skew for three key responses: 1) population persistence, 2) lag between range displacement and climate displacement, 3) location of maximum population sensitivity. Surprisingly, under our model assumptions, the direction of performance curve asymmetry does not strongly contribute to either persistence or lags. Conservation measures to support range-shifting populations may have most benefit near their environmental optimum or where the environmental dependence is shallow, irrespective of whether this is the ‘leading’ or ‘trailing’ edge. A metapopulation simulation corroborates our results.

## 1 Introduction

Climate change is driving range shifts of species across the world (Parmesan & Yohe 2003; Lenoir et al. 2020) making understanding the underlying ecological dynamics and drivers a key component of conservation efforts (Urban et al. 2016). For species with narrow environmental niches, their likelihood of survival will be determined by their relative rates of extirpation from sites at trailing edges and colonisation at leading edges (Kerr 2020). Range shifts are not instantaneous, creating communities not at equilibrium with their climatic niche (Svenning & Sandel 2013; Alexander et al. 2018; Rumpf et al. 2019; Lenoir et al. 2020). While species-distribution models (Elith & Leathwick 2009) can indicate the potential end state after climate change, dynamic models are required to examine how a species’ range may shift through time (Zurell et al. 2009, Alexander et al. 2017). To meet this need, many modelling approaches have been applied, spanning the full breadth of possible trade-offs between model precision, realism and specificity (Levins 1966).

Large generalised spatially-explicit simulation models (Brooker et al. 2007; Urban et al. 2012; Lurgi et al. 2015; Thompson & Gonzalez 2017; Thompson & Fronhofer 2019) are able to flexibly capture many processes and have been highly informative. However, the complexity of these models makes systematic interrogation of conclusions drawn from particular parameter choices a challenge. Results can depend strongly on the underlying assumptions of the model (Zurell et al. 2016). Moving-habitat integro-differential equation models (Zhou & Kot 2011; Kot & Phillips 2015; Harsch et al. 2017; Hurford et al. 2019), which include a continuous spatial element and discrete time, can capture many nuances of dispersal processes and are somewhat analytically tractable (Kot & Phillips 2015). However, conclusions are still largely restricted to inspection of simulation results.

To complement these more specific models, there remains a strong need for fundamental theory to identify critical determinants of climate change responses. Purely analytic models provide another perspective to the problem, providing an ‘all-else-equal’ baseline for consideration. To address this need, partial differential equation models building on well-established reaction-diffusion equation models (Cantrell & Cosner 2003) for invasive species and gene spread (Fisher 1937; Skellam 1951; Hastings et al. 2005), have been applied to climate change scenarios (Pease et al. 1989; Potapov & Lewis 2004; Berestycki et al. 2009; Li et al. 2014). This work has shown the critical rate of climate change that a species can survive to be a function of dispersal rate and population growth rate from rare, allowing direct comparison with data (Leroux et al. 2013).

However, to date, analytic models have been restricted to simple representations of species performance across environments. In particular, the functional dependence of the species’ intrinsic growth rate on its environment, i.e. the environmental performance curve (EPC), is assumed to be symmetric. It is widely appreciated that most species show highly asymmetric environmental responses (Savage et al. 2004), for example a gradual increase up to an optimum followed by a sharp decline. Disparities in environmental sensitivity between trailing and leading range edges may be expected to influence range-shift dynamics. For example, using a numerical approach, Hurford et al. (2019) demonstrated a case where the EPC asymmetry and dispersal interact to cause divergent effects depending on the direction of the asymmetry. Further, these asymmetries may be particularly relevant when underlying climatic variability is considered (Nadeau et al. 2017).

Here, we extend previous analytic theory to incorporate asymmetric environmental dependence and directly investigate the impact of asymmetry on three key responses: likelihood of persistence, range-shift lag and the location of peak sensitivity to conservation interventions. We do this by re-formulating the species-movement problem as a Schrödinger equation, unlocking mathematical results from the quantum mechanics literature. We then corroborate these results in a simulation exhibiting complex dynamics.

## 2 Analytic Theory

### 2.1 Setting and specification of core model

For simplicity we only outline the mathematical derivations here. Full details are given in SI 1 and SI 2 is an evaluated Mathematica notebook to replicate our analysis. Throughout, we assume a single spatial dimension (*x*) that spans a linear gradient in an environmental variable *E*, where initially *E* = *x*. Our *E* could represent any environmental variable, but is most easily conceptualised as an average temperature. Climate change is introduced by a linear time dependence of E: *E_x,t_* = *x* – *vt*, where *v* is the rate of change in *E*. Hence, the location where *E* = 0, moves with velocity *v*. In a warming scenario, *x* could be distance from a Pole or to a mountain peak and *v* would be negative. The correspondence between space and *E* means that *v* can also be interpreted as a ‘climate velocity’ of dimensions of Length × Time^−1^ (Brito-Morales et al. 2018).

A species’ local population growth rate, given the environment, is defined by an environmental performance curve (EPC) function *g*(*E*). Following e.g. Fisher (1937), Pease et al. (1989) and Hastings et al. (2005), we model changes in the local population density *b* = *b*(*x, t*) of a species through time t at location *x*, including dispersal at a rate *D*, as

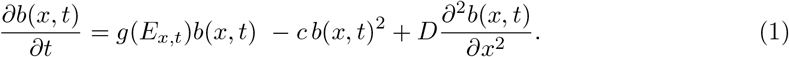

The effect of migration is incorporated via the final term of Eq. (1). The rate of net migration depends on the curvature of the population density to either side of the focal point. Populations near the peak of the biomass distribution (where the curvature is negative) lose population density to net-migration, while the edges of the distribution (where the curvature is positive) gain. The dispersal rate *D* is 1/2 times the mean squared displacement along *x* of a lineage per unit time and can related directly to the mean movement of individuals per generation (Kareiva & Shigesada 1983).

Central to our approach is a change of reference frame to track the region of suitable environment across space. We define *y* = *x* – *vt* and use *y* as our principal spatial variable (Fig. 1). This allows us to examine the distribution *u*(*y,t*) = *u*(*x* – *vt*, *t*) = *b*(*x, t*) of the population in a reference frame co-moving with the environment. Evaluation of the derivatives results in the following partial differential equation for *u*(*y, t*), equivalent to the equation for *b*(*x, t*) above:

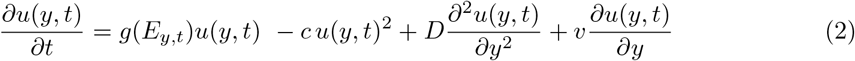

**Figure 1:**
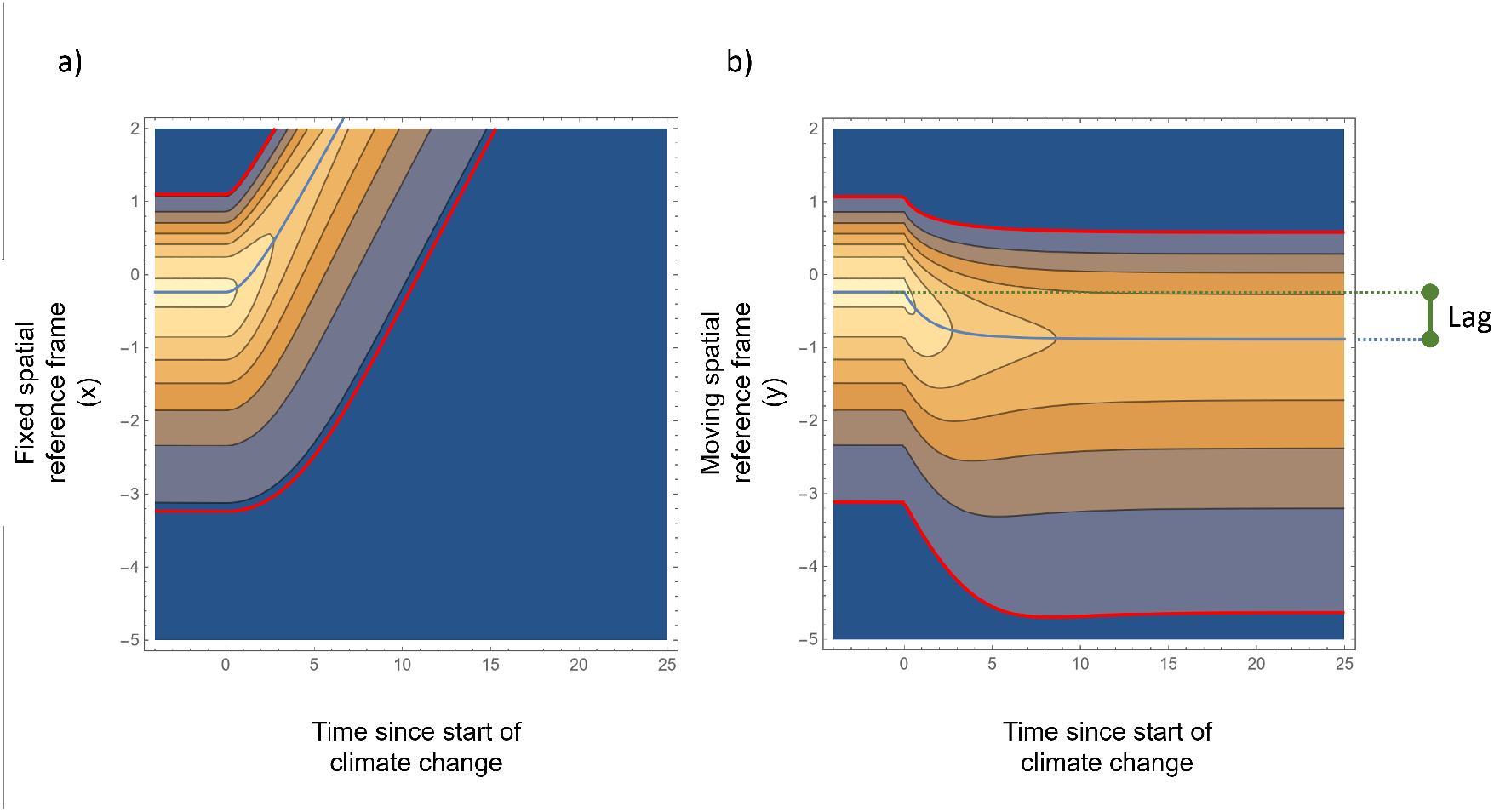
a) Example of movement of species range with climate change under a fixed spatial reference frame (*x*). Climate change rate (*v*) is set to +0.45 starting at t=0. Brightness of colour signifies population density, peak density is indicated by the central blue line. b) Species range through time under climate change in the moving reference frame (*y*) - parameters are otherwise identical to a). With ongoing climate change, after a period of adjustment the population reaches a steady travelling wave state, with a lower overall population density and a constant lag behind the moving climate. Parameter values are listed in the Mathematica supplement.

**Table 1:** Summary of parameters and variables used in the analytic theory.

|  | Term | Meaning |
| --- | --- | --- |
| *Core Model parameters* | $x$ | Spatial coordinate |
| | $v$ | Rate of climate change |
| | $y$ | Spatial location, reference frame moving with climate change $y = x - vt$ |
| | $b(x, t)$ | Population density at point $x$ and time $t$ (fixed reference) |
| | $u(y, t)$ | Population density at point $y$ and time $t$ (moving reference) |
| | $E$ | Environmental variable |
| | $g(E)$ | Environmental performance curve (EPC) function describing local population growth rate given $E$ |
| | $\psi(y)$ | Population distribution without climate change. |
| | $\lambda$ | Population growth rate |
| | $\lambda_0$ | Population growth rate without climate change |
| | $D$ | Dispersal rate |
| | $c$ | Strength of density dependence |
| *EPC parameters* | $w$ | Niche width parameter |
| | $a$ | Asymmetry of EPC |
| | $\phi$ | Environmental optimum, peak of $g()$ |
| | $R_{max}$ | Growth rate at optimum |
| | $\sigma$ | Standard deviation of temporal environmental variation |
| *Derived Values* | $v^*$ | Critical rate of climate change at which the population cannot persist ($\lambda = 0$) |
| | $\Delta^{space}$ | Lag in space between total climate displacement and displacement of population at steady state |
| | $\Delta^{time}$ | Lag in time between changes in the environmental values and the population density |
| | $S_{max}$ | Location of peak sensitivity of $\lambda$, relative to optimum |

### 2.2 Approximate model general solution using a Schrödinger Equation

For vulnerable populations - that is, populations that are close to extirpation - an approximate solution of Eqs. (1) and (2) can be constructed if the spatial distribution of the population prior to climate change (*v* = 0) is either known or has been computed from either Eqs. (1) or (2) (SI 1). We denote this spatial distribution with a function *ψ*(*y*), that describes the *shape* of the population distribution across space.

By focusing on vulnerable species the problem simplifies, because when populations are low the quadratic terms in Eq. (2) describing density-dependencies will generally be small compared to both the dispersal terms and the term containing the environmental performance curve *g*(·), with the latter two mostly balancing each other (S1.3). The effect of density dependence can then be taken into account as a small perturbation. As a result, the equilibrium solution *u*(*y, t*) will, up to a scaling factor, be very similar to the form that *u*(*y, t*) would attain while the population was growing from initial low abundance.

Mathematically, we therefore define *ψ*(*y*) not directly as the solution of Eq. (2) for *v* = 0 but as the solution of the corresponding equation without density dependence for a population growing from low abundance with the maximum feasible growth rate λ_0_:

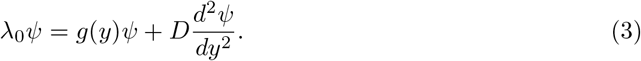

This equation defines *ψ*(*y*) up to a constant factor that does not matter for the following.

We show in S1.2 and S1.3 that from *ψ*(*y*) and λ_0_, an approximation of the species’ response to climate change at any given velocity *v* can be obtained without solving another differential equation. Instead, the populations’ approximate distribution in the presence of climate change (with velocity *v*) is

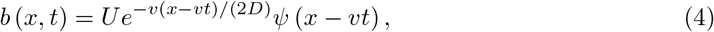

where *U* is a scaling parameter given by

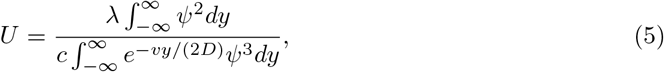

and

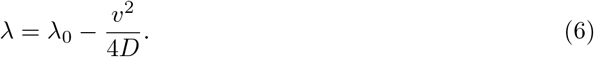

Under our assumptions, Eq (4) holds up to a relative error of the magnitude of λ/ max_*y*_ *g*(*y*). The population’s actual distribution prior to climate change is a good first approximation of *ψ*(*y*) (= *ψ*(*x*)) and can be used in place of *ψ* above. Alternatively, Eq. (3) can be solved numerically. Here, however, we make use of another possibility. This draws on the formal equivalence between Eq. (3) and the time-independent Schrödinger equation (Schrödinger, 1926) for the wave function *ψ* of a particle moving in a one-dimensional energy potential *V*(*y*):

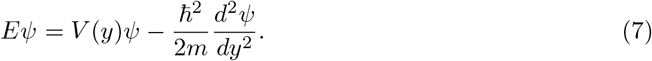

In this equation, *E* denotes the total energy, *m* is the particle’s mass and 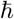 is a constant. Comparing terms, one sees that the potential energy *V*(*y*) corresponds to the environmental performance function *g*(·) (with a sign flip). The final term describes the kinetic energy of the particle and corresponds to the dispersal term in Eq. (3). The value of these correspondences come from the fact that Eq. (7) has been studied extensively in quantum physics. This has led to a strong body of intuition about the nature of the solutions of the eigenvalue problem given by Eq. (7) and the discovery of many functional forms for *V*(*y*) for which closed-form solutions can be derived (Mattis 1993).

Importantly, we are not adopting an interpretation of population density as a quantum mechanical wave function. Our results are distinct to the fundamental uncertainty aspects of quantum mechanics that may be familiar to some readers, although previous authors have suggested that those could have ecological applications too (Bull 2015; Real et al. 2017). Rather, we are transferring existing mathematical results from the quantum mechanics literature. We only consider real valued-solutions and so our problem is closely related to classical reaction-diffusion models (Nagasawa 1993).

### 2.3 Representing Environmental Performance Curves

In our model the function *g*(*E*) describes how the local intrinsic growth rate of a population (i.e. without dispersal effects) depends on environmental conditions. It could be any function that is negative as *E* becomes very large or very small. However, by selecting one of the many functions for which Eq. (7) has been solved analytically (for an accessible list see http://wikipedia.org/wiki/List_of_quantum-mechanical_systems_with_analytical_solutions)considerable progress can be made.

An asymmetric environmental performance curve (Fig 2b,c) can be defined in analogy to the Morse potential (Morse 1929) function

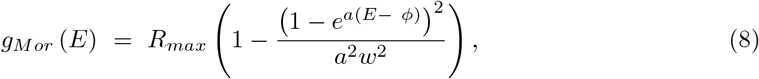

where *R_max_* is the local intrinsic growth rate of the species at the environmental optimum *E* = *ϕ, w* is a niche width parameter and *a* is a non-zero asymmetry parameter. For simplicity we will assume the environmental optimum to be at *ϕ* = 0 throughout. We assume that |*aw*| < 1 throughout, which assures that both very high and very low values of *E* lead to negative growth rates *g_M or_* (*E*). EPC *g*(*E*) provided in other functional forms are best approximated by *g_M or_* (*E*) by matching the first three derivatives at the point *ϕ* of maximum performance (*g*′(*ϕ*) = 0). This is achieved by setting *w* = (2*R_max_*/|*g*″(*ϕ*)|)^1/2^ and *a* = *g*′′′ (*ϕ*) /3*g*″ (*ϕ*).

**Figure 2:**
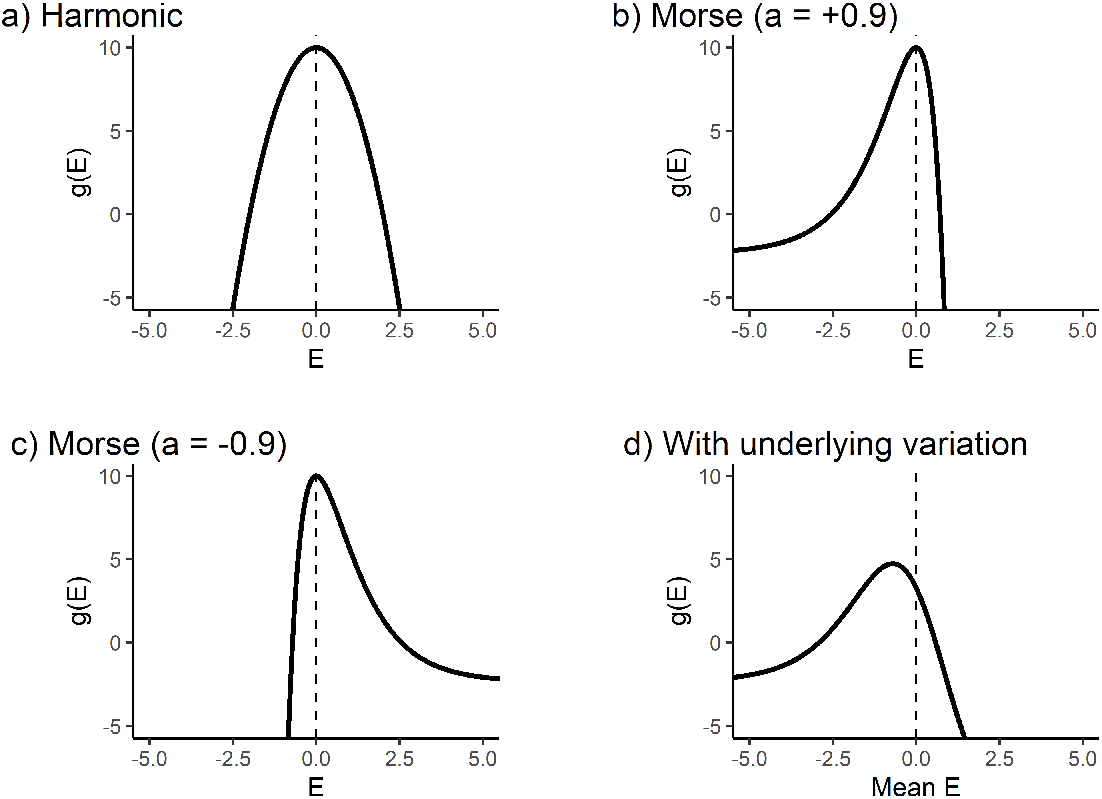
Illustration of environmental performance curves under different models that permit analytic solutions. In each case the species has an optimum *ϕ* = 0 and *R_max_* = 10 a) harmonic potential function, *w* = 2. b) Morse potential function, where *a* = +0.9, *w* = 1 c) Morse potential function *a* = –0.9, w = 1, d) Morse potential function as ‘*g*(*E*)’, but incorporating the effects of a variable *E*, (mean 0, *σ* = 1, *a* = +0.9, *w* = 1). Note how with the peak of the curve has shifted with the introduction of this climate variability.

Care is needed when referring to skew direction - a *positive* value of *a* leads to a ‘negative’ or left-tailed skew (Fig 2b,c) in terms of *E*. Our *E* variable declines with climate change - if *E* is a temperature variable, lower values are therefore warmer, and positive *a* values would be described as ‘warm-skewed’ because the heavy tail is to the warmer (more equatorial, lower elevation) side of the optimum (Hurford et al. 2019).

On top of these underlying functions, the effect of high-frequency temporal environmental variation can be modelled by convolving these curves with a probability density function that represents the environmental variation. When environmental variability is represented by a Gaussian distribution with standard deviation *σ*, the convolution with the Morse potential function Eq (8) maps onto a transformed Morse potential. In Fig 2d, we show the effect of introducing variability with *σ* = 1 on the overall effective environmental performance curve: it softens the edges and shifts and flattens the peak of the performance curve (Ruel and Ayers 1999).

## 3 Analytic Results

We use our model to derive analytic predictions for how three response properties depend on key parameters – 1) the capacity for species to sustain themselves under climate change, 2) the lag between a species’ distribution and where it ‘should’ be if it kept pace with climate change, and 3) the location where conservation interventions would be most efficacious.

### 3.1 Response 1: Critical speed of climate change

The sign of the low-density growth rate λ under climate change is crucial for a moving population (Grainger et al. 2019). If positive, a population at low abundance will grow (despite climate change), to the equilibrium state described by Eq. (4). If negative, the population will to decline to extinction. By Eq. (6), climate change is always detrimental to fitness. Remarkably, this fitness decline is entirely independent of the form of the EPC *g*(*E*). Also of interest is that the impact of climate change rate on population fitness is quadratic, implying that linear extrapolations from observations of slow change will not correctly predict the impact of faster changes.

Equation (6) can be rearranged to determine the critical speed of climate change (*v*\*), at which a species can no longer keep up and will go extinct (i.e. λ = 0):

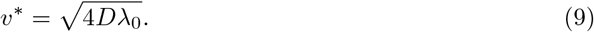

Alternatively, one can solve for *D* to identify the critical rate of dispersal that a species must exceed to maintain its population (Hastings et al. 2005; Leroux et al. 2013). Again, neither of these results depend on the shape of *g*().

The shape of the performance curve does, however, influence the pre-climate change population growth rate (λ_0_). Using the asymmetric EPC defined above, we can obtain (S2.3) a relatively simple expression for the intrinsic population growth rate,

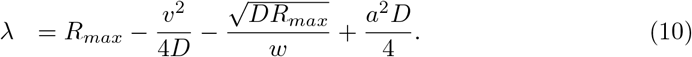

This is valid as long as *D* < 4*R_max_a*^−4^*w*^−2^ (S1.6).

The first three terms of Eq. (10) correspond to previously found conclusions about the impact of climate change in the case of a symmetric (quadratic) EPC (Pease et al. 1989). Consistent with intuition, the model predicts that greater maximum population growth rate increases fitness and that climate change is detrimental to population fitness. The effect of dispersal rate *D* is multifaceted – the second term shows how greater dispersal can offset the impact of faster climate change, but the third term shows a negative effect of greater dispersal due to losses from the central part of the population. This effect is mitigated by larger niche widths.

The fourth term predicts that asymmetry (*a*) acts quadratically, and is therefore independent of skew direction – whether the long tail in performance is on the leading or trailing range edge is not relevant to population fitness. This follows naturally from the result that asymmetry only influences the baseline population fitness λ_0_, which does not have any sense of directionality. Grouping the terms can identify λ_0_ = *R_max_* – (*DR_max_*)^1/2^*w*^−1^ + *a*^2^*D*/4, i.e. most of the parameters in Eq. (10) contribute only to the λ_0_ component, not the response to climate change as such.

Similarly, high-frequency temporal variation impacts only the form of *g*(), and so can be shown not to alter the marginal impact of climate change in our model (Eqs. (6), (9)). That said, *σ* does impact the overall expression for λ in a complex manner (see S2.3.2). The marginal effect of increasing *σ* from a low value on λ,

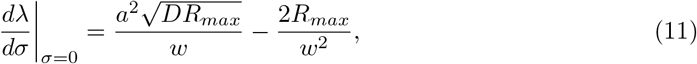

is easier to interpret. Taking into account the condition *D* < 4*R_max_a*^−4^*w*^−2^ for Eq. (10), increasing variability will always lead to a reduction in growth rate. However, it shows that the marginal effect of temporal variation on fitness is dependent on a large number of model terms. The range width denominators confirm the intuition that temporal variation is most influential with narrow ranged species. Both dispersal and asymmetry in Eq. (8) effectively widen the niche and correspondingly reduce the negative impacts of environmental variation. Again, the direction (sign) of the asymmetry is not relevant since *a* enters quadratically.

### 3.2 Response 2: Range shift lag behind climate change

Since both dispersal and population growth take time, there is expected to be a measurable lag (in space and time) between the suitable climatic range and the distribution of a population (Alexander et al. 2018). With a constant rate of climate change, and assuming a species’ population is able to sustain itself, it will eventually reach an equilibrium in coordinates co-moving with the changing climate. Even in an idealised model system like ours, there are multiple metrics that can be used to describe a species range in space (Yalcin & Leroux 2017). Here we measure lag in the moving reference frame between the pre-climate change peak of population density and the peak with climate change once it reaches the travelling wave stage (Fig 1).

Using the asymmetric EPC including the effect of temporal variation, we can derive expressions for these lags (S2). Focusing on the case where the rate of climate change *v* is relatively small (*v* → 0), the steady-state lag (Δ) in time becomes:

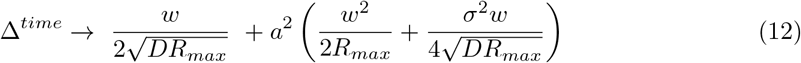

This lag in time can be converted to lag in space by multiplying by the velocity of climate change. If ‘lag’ is instead measured in terms of the centre of mass of the population’s distribution, identical results are obtained to lowest order in *v* and *a* (S2.4). Where there is no asymmetry (*a* = 0), only the first term is relevant. The denominator reaffirms the intuition that lags are reduced by greater dispersal and larger maximum growth rates. The numerator shows that lags are greater with wider niche widths – the population is not forced to move as rapidly when a larger part of the environment is suitable.

From the second term of Eq. 12 it can be seen that the exact impact of *a* depends in a complex way on other parameters but responds only to the magnitude of asymmetry, not the sign. Overall, greater maximum growth rate *R_max_* and greater dispersal *D* always decrease that lag and greater niche width *w*, asymmetry *a*, and climate variability always enhance it. The effect of climate variation *σ* is tied to the asymmetry of the EPC, with variation combining with asymmetry increasing effective niche width, while in the symmetric case *σ* has no effect.

### 3.3 Response 3: Sensitivity to Conservation Actions

Our model can be analysed to examine where interventions are most consequential for the overall population growth rate, and might therefore be of particular conservation priority. We do this by assessing the sensitivity of the overall population λ to local perturbations in growth rate across space. Note that we use the moving spatial reference frame *y* (and optimum *ϕ* = 0), so that locations are relative to the environmental optimum at each point in time as climate change progresses. This method is analogous to determining the most sensitive life-stage (e.g. Caswell 2012, 2019), but substituting age by space. The sensitivity of λ is proportional to the product of the reproductive value and the abundance at each environment in the stable distribution.

Computing the shift of the peak sensitivity *S_max_* relative to the value of *y* where the environment is optimal, we find

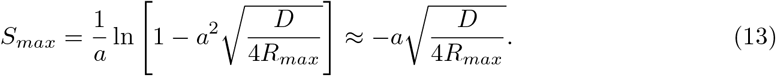

Remarkably, the shift *S_max_* away from the optimum environment depends neither on the rate nor direction of environmental change. In S1.5, we show that this independence on *v* is a highly general result valid for any functional form of the EPC. For symmetric EPC, there is no shift at all, but for our asymmetric EPC maximum sensitivity always occurs on the long-tail side of the environmental optimum (Figure 3).

**Figure 3:**
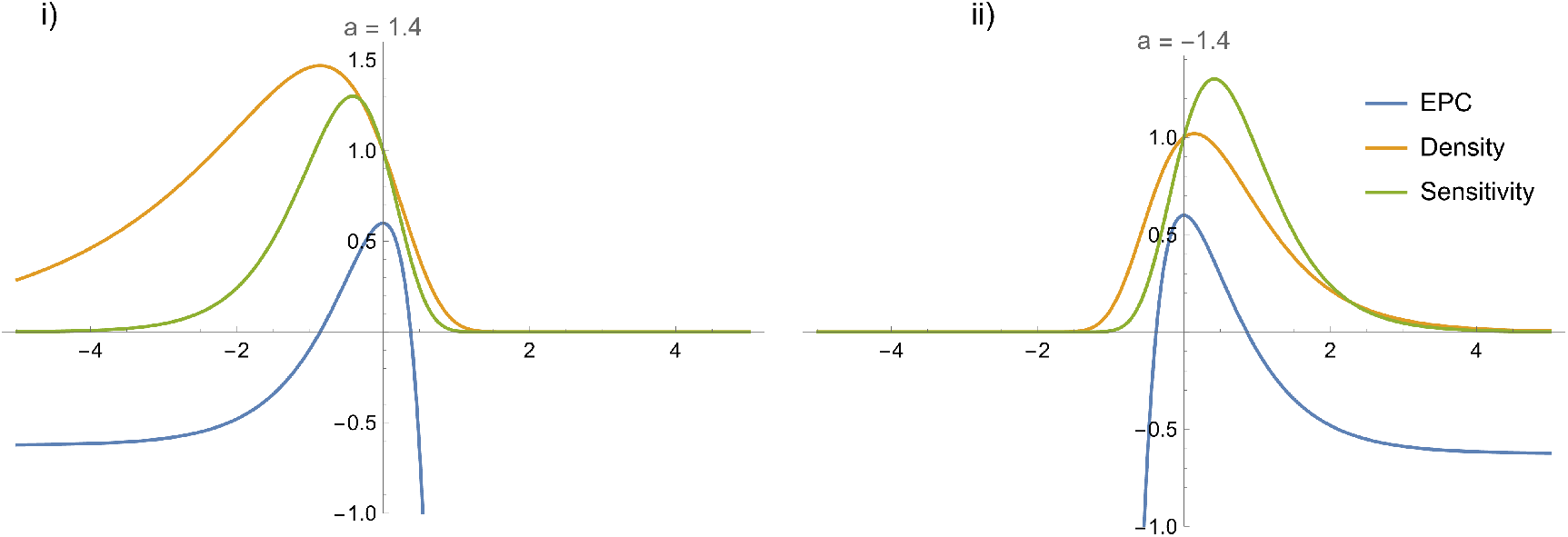
Distribution of sensitivity to intervention and population density at equilibrium with climate change. In both panels, the horizontal axis is in *y*, the species leading edge is to the right, and the trailing edge to the left. Apart from the asymmetry of the EPC, parameters are the same, only the direction of asymmetry varies. Horizontal axis = *y* (the moving reference frame) where 0 = *ϕ*, the species environmental optimum. Three quantities are plotted: the EPC function, showing environmental optima at *y* = 0 (blue), the population density (orange, *ψ*, standardised to equal 1 at *y* = 0), and the sensitivity to intervention (green, also scaled)

## 4 Comparison with simulations

To derive analytic results, our framework makes a number of simplifying assumptions. To test the translatability of our predictions to more complex systems, we built a spatially-explicit population model. Expanding on the continuous-time and continuous space analytic results, we built this model to include randomness and discreteness in both space and time. Because of the inherent challenges in estimating key parameters we focus on qualitative comparisons between our analytic predictions and the simulations, but show in SI 4 that quantitative predictions are of the correct order of magnitude. Where possible we use the same symbols for simulation and analytic parameters where they are similar, but note that they are not directly comparable because of differences in overall model structure.

### 4.1 Model Specification

Full details of our simulation model are given in SI 3. Briefly, we generated rectangular arenas with dimensions 40×10, each with 200 randomly located nodes connected to their nearest neighbours (Figure 4a). A single environmental variable *E* linearly increases along the *x*-axis of each arena from 0 to 40. Population dynamics of each species were modelled in discrete time and followed logistic growth with dispersal between nodes, where the intrinsic growth rate *R_max_* depended on the current environment at that node through performance functions based on Eq (8), with *R_max_* = 10, *w* =1 and *a* = ±0.9). To represent temporal environmental variability, for each set of 15 timesteps (which we refer to as a ‘year’) we added a value drawn from a Gaussian distribution (mean=0, *σ* = 0.5) to all *E* values during that year. Immigration into neighbouring sites was determined by an exponential distance decay function, with dispersal rate chosen such that extinctions are possible once climate change is introduced.

**Figure 4:**
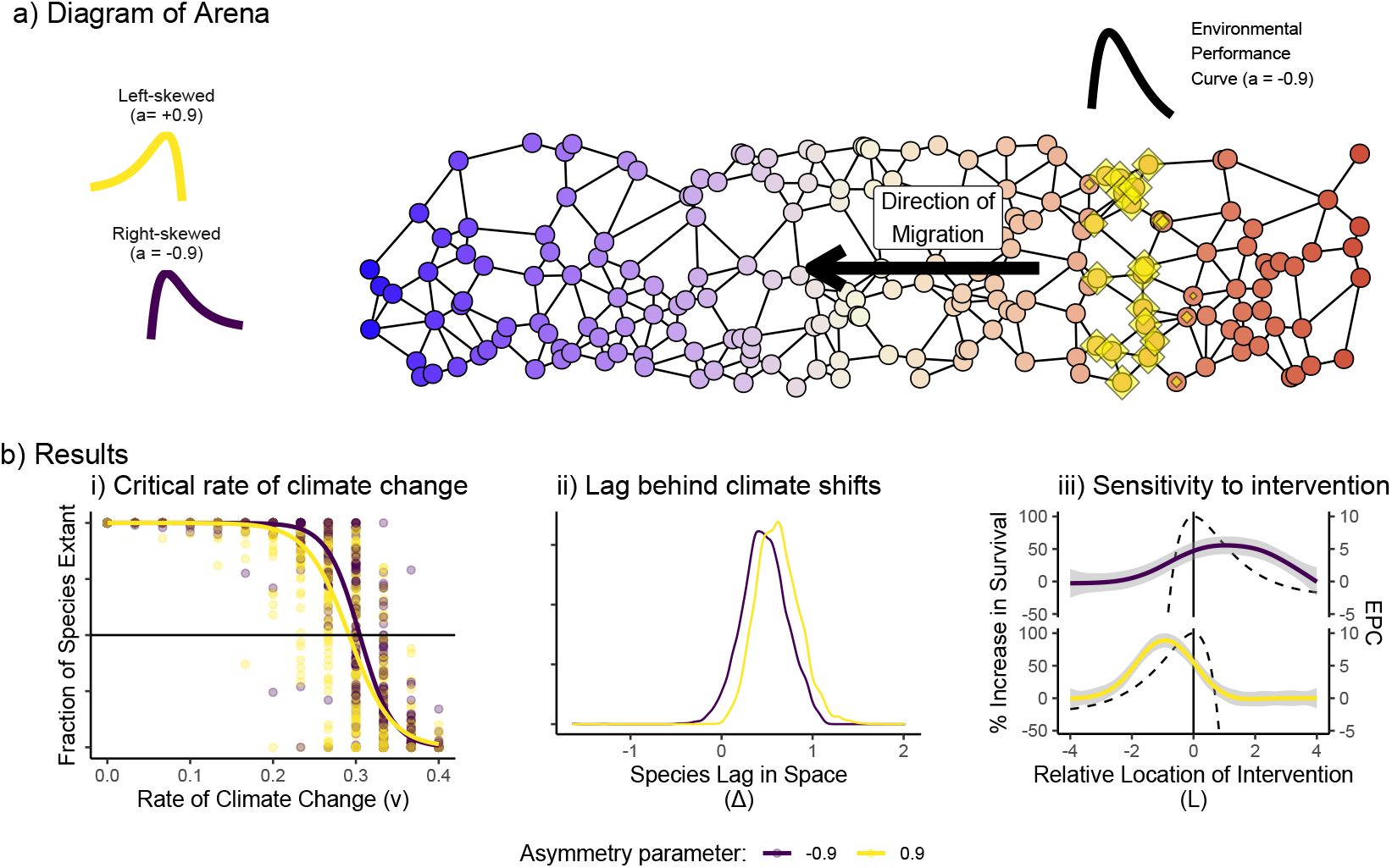
Metapopulation simulations setting and results. a) Illustration of example virtual region of connected habitats the species inhabit in the simulation. Initial mean environmental values (*E*) are given by the blue (low) to red (high) colouration. The size of the yellow diamonds illustrates the population density at each site. The environmental performance curve of the species (right-skewed, *a* = –0.9) is drawn above. With climate change and an increasing underlying environmental variable, the species will have to shift its range leftwards. b) Numerical results from the simulations, partitioned and coloured by the direction of the skew of the EPC. i) The speed of climate change (*v*) at which the proportion of species that are able to survive falls below 0.5. Lines are fitted binomial GLMs. ii) Density plots of the lag by which species fall behind the movement of climate at a moderate rate of climate change. iii) Observed increase in survival during climate change, at different locations of conservation intervention relative to each species’ optimum. Solid coloured lines are GAMs fit through simulation results with 95% confidence intervals. Black dashed lines illustrate the environmental performance curves for reference. Peak efficacy aligns with the long tail of the environmental performance curve in both cases. For all three responses, the qualitative predictions of our analytic model are supported.

We assembled 100 sets of species with either direction of environmental performance skew (*a* = +0.9 or −0.9). We did not test a symmetric environmental performance curve case, as it is not possible to standardise all aspects of the performance curve for a fair comparison. For each set, 100 species were generated with environmental optima (*ϕ*) each drawn from a uniform distribution between 20 and 30, maintaining a region of suitability throughout the simulation to mitigate possible edge effects. The arenas were seeded with initial colonists and the model integrated for 200 ‘years’ to fill their initial range. Species that at any point fell below a threshold biomass (10^−6^) across all nodes were considered extinct and removed.

We examined three response metrics that could be derived from both the simulation model and the analytic model, to assess if the pattern of parameter dependencies holds.

### 4.2 Critical rate of climate change

We sequentially simulated each assembled set of species under varying rates of climate chance (*v* = 0 to 0.5 in steps of 0.05) and identified the fraction of species that had fallen below 1% of their starting total population size at the end of 50 ‘years’ of climate change. In line with our expectations, there was a relatively precipitous decline in survival chance. Both *a* = +0.9 and −0.9 showed very similar responses (Figure 4bi). We confirmed that the asymmetry parameter was indeed impacting *v\** with trials of *a* = −0.7, 0.3, 0.3 and 0.7 (SI 3). We fit a generalised linear mixed-effects model that included as main predictors *v, a*, their interactions and assemblage as a random effect to estimate *v*_50_, the *v* at which the 50% of species go extinct. When left-skewed (*a* = +0.9), *v*_50_ = 0.292, while when right-skewed (*a* = −0.9), *v*_50_ = 0.305.

### 4.3 Movement Lags

We tested the lag in movement of the centre of mass of the population distribution for each of the assemblies during climate change. We first assessed the starting location of the ‘centre of mass’ of each species before any climate change (*m_i,start_*) as the average *x*-coordinate of each node, weighted by the biomass of species *i* at each node, averaged over 20 years. We then ran 70 years of climate change at *v* = 0.1, and measured the average lag in space 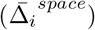 for each species over the final 20 years:

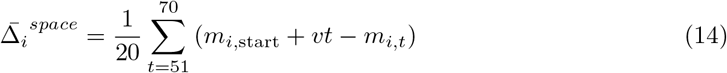

Overall, *a* = +0.9 led to a marginally greater average spatial lag (0.618) than *a* = −0.9 (0.467), but this was small compared to the overall amount of variation in the simulations (Fig. 4bii).

### 4.4 Location of Peak Sensitivity

Localised conservation interventions were represented by increasing the intrinsic growth rate by 2 at all sites within an intervention window 1 spatial unit wide. This band was centred *L* spatial units in the *x*-dimension from the optimum of each species. Climate change was introduced at a rate of *v* = 0.28, which in the absence of intervention would cause around half the species to go extinct. When climate change was introduced, the intervention window moved with it. The simulation of 50 years of climate change was repeated with values of *L* ranging from +3 to −3 (in steps of 0.5) and the percentage increase in species surviving the climate change period with the conservation intervention, compared to simulations without the intervention was recorded.

In line with the analytic expectations, we found that the location where the sensitivity had the most impact depended on the direction of the skew of the asymmetry of the EPC (Fig. 4biii). Interventions were most efficacious at preventing extinctions when they were located away from the optimum on the shallow environmental sensitivity (long tail of the EPC) side of the moving range, regardless of whether this was the ‘leading’ or ‘trailing’ edge of the range.

## 5 Discussion

Our results shed new light on how the shape of a species environmental niche and other key drivers may impact responses to climate change. Our finding that while the extent of asymmetry is potentially highly influential, the *direction* of skew is not relevant to either the likelihood of population persistence or the range shift lag in our model is particularly surprising. Metaanalyses of the rates of species range shifts have not identified strong trait or taxonomic signals, instead identifying range size and habitat breadth as the most informative predictors (Maclean & Beissinger 2017; Lenoir et al. 2020). If it is assumed that the direction of environmental performance asymmetry is linked to taxonomy, this would align with our results, however data specifying environmental performance curves is not presently available to conduct strong tests.

In physics, beyond the simplest case of the movement of a hydrogen atom in a vacuum, the complexities of multi-body problems limit the resolution with which predictions can be made. The same is true in ecology, and there are a large number of additional processes including species interactions, evolution, demography and environmental heterogeneity that will influence the observed dynamics (Urban et al. 2016). Analytic calculations will struggle to concurrently include the full diversity of processes that simulation models can achieve, and certain features pose particular challenges, for example heavy-tailed dispersal models that do not have moment-generating functions (Liu & Kot 2019). Nonetheless, the expansion of the analytic theory is fundamental to building a foundation of expectations for how the natural world will react.

Using our model, we demonstrated that interventions will have the greatest benefit on a rare range shifting species when located somewhat near the centre of the range, in the region corresponding to the long-tail of an asymmetric performance response. The result is consistent regardless of whether this region is towards the leading or trailing edge of the at-risk species’ range. This conclusion is somewhat different to discussions about whether it is most helpful to support a species at its trailing edge (where it is at risk of imminent disappearance) or towards its leading edge (where assisted translocation is possible) and could form a possible rule-of-thumb when information is sparse.

### 5.1 Applicability and Scope

In principle, the parameters in the analytic model could be directly measured for natural populations (e.g. Leroux et al. 2013). However, measuring dispersal rates remains challenging, especially given the importance of rare long distance events (Kerr 2020). The predictability of dispersal and colonisation rate is limited even under tightly controlled experimental conditions (Melbourne & Hastings 2009) leading to hard limits to the accuracy of any direct model prediction.

In our analysis, we interpreted *b*() as denoting a population density (individuals per unit area). However, one can instead interpret *b*() as the density of patches occupied (occupied patches per unit area), in the spirit of metapopulation modelling introduced by Levins (1969). This second interpretation may lead to a closer alignment between the assumptions of the model and empirical realities, as well as being more directly amenable to empirical verification or parameterisation using grid-cell occupancy data.

An important step in the derivation of our analytic results is the focus on species that are close to their extinction threshold. This allows the impact of density dependence to be assumed minimal and the application of perturbation theory. However, a steady state of a system such as Eq. (2) can broadly be achieved in two ways - either through high dispersal and low density-dependence, or through higher density-dependence and much lower influence of dispersal on growth rate. We examined species with minimal density-dependence, and therefore dispersal terms will strongly influence local growth rates, including a not-insignificant reduction in population density in central areas due to emmigration. Our model is therefore likely most directly applicable at local scales where mass effects are more significant than occasional rare long-distance dispersal events.

This context is helpful to understand why our findings contrast with a previous modelling result. Hurford et al. (2019) found through simulations of an integro-differential equation model that positively skewed EPCs were associated with reduced lags compared to negatively skewed EPCs. Hurford et al. attributed this result to increased immigration into newly suitable areas from high-density populations near the leading edge. A key difference between our assumptions that may explain the discordance is that we focus on species close to an extinction threshold, while in Hurford et al.’s model extinctions do not occur.

### 5.2 Limitations and extensions

We base our results around the properties of a stable travelling wave and use the toolbox of quantum physics to precisely describe its motion. Our analytic framework is therefore orientated around long-term equilibrium solutions. Yet, there is considerable scope for the analysis of transient dynamics following the start of the perturbation (in this case climate change), which may be more representative of available ecological observations (Hastings 2016). While our work significantly extends the scope of analytic theory in this field, there remain many further processes that are not directly considered. Our approach to modelling temporal variability effectively only modifies the underlying growth curve. It therefore only indirectly captures the impact of discrete stochastic events, that can be highly influential over short time frames relevant to contemporary climate change responses.

Further, neither our analytic model nor simulations explicitly include interactions between species precluding potential ‘box-car effects’, where competitive interactions slow the rate of advance (Urban et al. 2012; Legault et al. 2020), or the potential for extirpation from areas due to the climate-driven arrival of new species. Although the shape of the environmental performance could be attributed to indirect biotic as well as direct influences of the environment, this would not necessarily capture the complexities of interaction with other species, particularly when climate variability is considered (Terry et al. 2022). Models for interacting species within the reaction-diffusion framework have been developed (Cantrell & Cosner 2003, Potapov and Lewis 2014) and incorporating interactions within a moving-environment framework is an interesting future avenue of research. Lastly, our model assumes fixed species traits, but species have the potential to adapt their traits to changing climates through plasticity or evolution (Hoffmann & Sgrò 2011). Trait adaptation can be built directly into the partial differential equation framework (Pease et al. 1989, Chevin et al. 2010) and represents a further promising area for further work.

### 5.3 Conclusion

Although spatial partial differential equation models have a long pedigree within ecology (Fisher 1937; Hastings et al. 2005), our results show how rich seams of results remain to be harnessed to generate fresh ecological perspectives and more detailed baseline expectations. Our focus on the special - but critically important - case of species close to extirpation allows a simplification to an essentially linear problem and the incorporation of knowledge from other disciplines that can bring new and surprising analytical insight.

## Acknowledgements

All authors were funded by NERC grant NE/T003510/1 (‘Mechanisms and prediction of large-scale ecological responses to environmental change’).

## Data accessibility statement

This paper uses no empirical data. All code used (R and Mathematica) and simulation results are publicly available at https://github.com/jcdterry/AnalyticRangeShift_Public and should the manuscript be accepted will be archived in an appropriate public repository (Zenodo).

## S1 Mathematical Derivations

We develop our analytic theory in two main steps. In the first, we formulate and analyse a partial differential equation (PDE) model for the distribution of a species’ population in a moving environmental gradient. Several results valid for essentially any environmental performance curve (EPC) are derived. In the second step, we derive more detailed analytic predictions for a specific choice of the functional form of the EPC.

### S1.1 Setting

We assume that the relevant environmental variable changes linearly along the *x* spatial axis, while being essentially constant in the other spatial directions. We denote by *b*(*x, t*) the time-dependent (*t*) distribution of a species’ population along the *x* axis. This can be understood in two different ways: (1) as a population density (individuals per unit length) or, (2) in the spirit of metapopulation modelling introduced by Levins (1969), as the density of patches occupied by the population near *x* (occupied patches per unit length). In developing the analytic theory, we will stick with the first, more conventional interpretation, but the second interpretation may be more appropriate in certain cases.

If the environmental gradient is not too steep, and so the range over which the performance curve permits a population to grow not too narrow, we can course-grain over individuals (1st interpretation) or patches (2nd interpretation) and model the dynamics of *b* = *b*(*x, t*) using the PDE model:

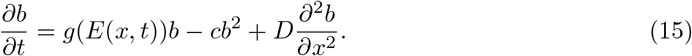

The first term on the right-hand-side describes population growth (*g*(*E*) > 0) or decline (*g*(*E*) < 0) dependent on the environmental variable *E* = *E*(*x, t*), the second term intraspecific competition with competition coefficient c, and the third term random dispersal of individuals, modelled as Fick diffusion.

We assume that the environmental variable *E* changes linearly in space and that the point in space where *E* = 0 moves along the *x* axis at a constant velocity *v*. Then *E* = *E*_0_ + *p*(*x* – *vt*) with some constant *E*_0_. We measure lengths in units such that *p* =1 and chose the origin of the *x* axes such that *E* = 0 at *x,t* = 0, implying *E*_0_ = 0. This simplifies Eq. (15) to

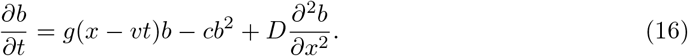

Next, we introduce a spatial coordinate co-moving with the environmental variable. We define *y* = *x* – *vt*, so that *x* = *y* + *vt*. Writing *b*(*x, t*) = *u*(*x* – *vt*, *t*) = *u*(*y, t*), we obtain for *u* = *u*(*y, t*) the equation:

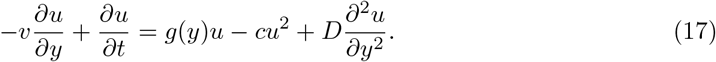

### S1.2 Invasion fitness

An ecologically important quantity is the invasion fitness of the species in question, i.e., its long-term population growth rate at low abundance. This can be obtained by dropping the quadratic term in Eq. (17) (reflecting an assumption that it is negligible because u is small, see next section) and looking for solution of the form *u*(*y, t*) = *e*^λ*t*^*u*_inv_(*y*) (with *u*_inv_(*y*) → 0 as *y* → ±∞), where λ is the invasion fitness. Mathematically, this leads to the eigenvalue problem

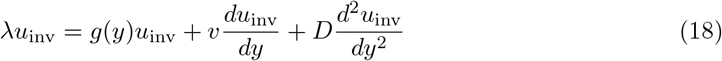

which can be written equivalently in Sturm-Liouville form

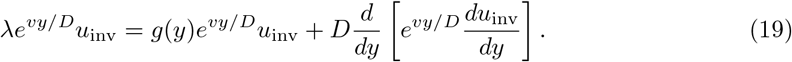

By Sturm-Liouville theory (Al-Gwaiz, 2008), there is, up to a constant factor, no more than one non-negative solution *u*_inv_(*y*) ≥ 0 and corresponding eigenvalue λ solving this problem. Below, in section S1.4, we shall study this solution for special cases in more detail, assuming that it exists.

### S1.3 Perturbation theory

First, however, we will use perturbation theoretical methods to obtain from solutions of Eq. (18) approximate solutions of Eq. (17) and insights about the responses of populations to environmental change and management interventions.

For simplicity, we consider only equilibrium solutions *u*(*y, t*) = *u*(*y*). Standard procedures (Chen et al. 1996) can be applied to extend this line of thought to time-dependent solutions.

Of particular interest is the situation where the focal species is at risk of extinction because λ is positive but close to zero. To study this case, we assume that *u* is everywhere so small that the quadratic term in Eq. (17) is small compared to the other terms (later results will vindicate this assumption). We introduce a book-keeping parameter *ϵ* to keep track of the order at which the small effect of the non-linearity contributes to corrections of the solution (we will set *ϵ* =1 in then end) this. To make the problem accessible to singular perturbation theory, we subtract λ*u* from the right hand side of Eq. (17) and then add *ϵ*λ*u*, with net zero effect. Then we write the time-independent form of Eq. (17) as

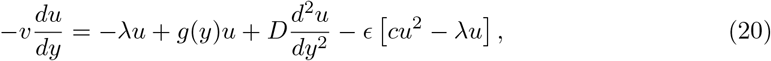

Following a standard procedure of perturbation theory, we decompose

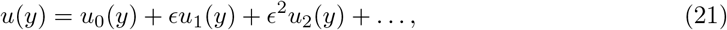

insert this expansion into Eq. (20), and sort terms by powers of *ϵ* (noting *ϵ*^0^ = 1):

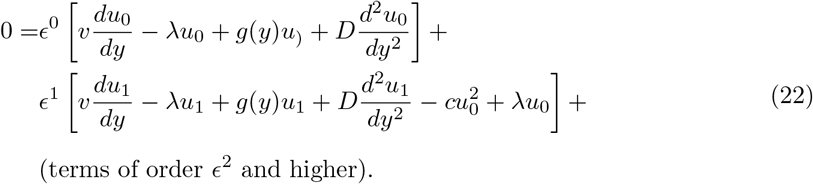

This equation is now solved at each order in *ϵ* separately. We have constructed this expansion such that at order *ϵ*^0^ the general solution is *u*_0_(*y*) = *U u*_inv_(*y*), with the constant *U* > 0 to be determined. Solving the equation for *u*_1_ at order *ϵ* is not always possible, because the linear operator *L* defined for arbitrary bounded and smooth functions *f*(*y*) as

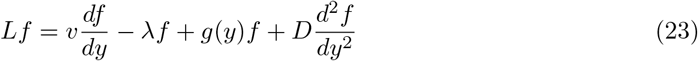

has a non-trivial null space *f* = *u*_inf_ and is therefore not invertible. According to Fredholm theory (Zeidler 1995), solvability requires that for functions *u*^+^(*y*) in the null space of the adjoint *L*^+^ of operator *L*, the condition

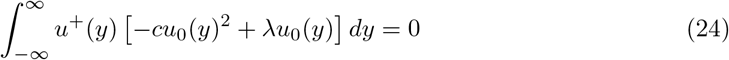

is satisfied. By standard arguments (Zeidler 1995), the adjoint operator *L*^+^ is obtained by flipping the sign of all *y*-derivatives *d/dy* in Eq. (23), giving

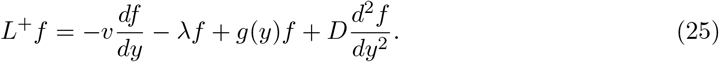

The null space of *L*^+^, that is, functions *u*^+^ satisfying *L*^+^ *u*^+^ = 0, are, up to a constant factor, of the form

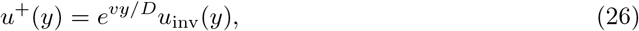

as is verified by direct evaluation. Ecologically, *u*^+^(*y*) provides, up to a constant factor, the reproductive value of individuals at location *y*. With this in mind, the factor *e^vy/D^* in Eq. (26) is ecologically plausible: as tendency, it assigns to individuals ahead in the range shift a higher reproductive value than to those lagging behind.

Putting Eq. (26) and *u*_0_ = *U u*_inv_ into the solvability condition Eq. (24), the condition can be evaluated further, yielding the hitherto unspecified constant

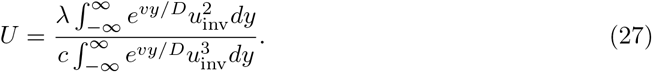

Since *u*_inv_ enters quadratically in the numerator and to third order in the denominator of *U*, Eq. (27) implies that *u*_0_ = *U u*_inv_ scales as λ/*c*, i.e. the magnitude of *u*_inv_ cancels out. It follows that, in order to balance contributions from *u*_0_ in *ϵ*^1^ term in Eq. (22), the term *g*(*y*)*u*_1_ must scale as λ*u*_0_. We can therefore estimate that the contribution *u*_1_ to *u* is by a factor of the order of magnitude of λ/max*_y_ g*(*y*) smaller than the contribution *u*_0_. As λ approaches zero from above, the term *ϵu*_1_(*u*) in Eq. (21) (with *ϵ* set to 1) therefore becomes negligible compared to the leading term. One can argue similarity for the higher-order corrections. Hence, populations in equilibrium but close to extirpation are approximately distributed with a density

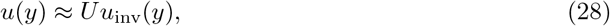

with *U* given by Eq. (27). (In the study of non-linear dissipative systems, Eq. (28) is known as the *weakly nonlinear approximation* (Stephenson & Wollkind 1995) of the equilibrium of Eq. (17).) This result establishes how the solution of the eigenvalue problem Eq. (18) essentially determines the equilibrium solution of the non-linear problem Eq. (17), especially for species close to extirpation.

### S1.4 Population fitness under climate change

We now study Eq. (18) in more detail. The equation can be simplified by writing *u*_inv_(*y*) as uinv(*y*) = *e*^−*vy*/(2*D*)^*ψ*(*y*) with a new unknown function *ψ*(*y*). Putting this into Eq. (18) yields an eigenvalue problem involving *ψ* = *ψ*(*y*):

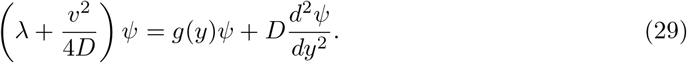

This formulation of the problem has the advantage that it depends on the velocity *v* of environmental change only on the left-hand side. To understand the significance of this result, let us assume that the species in question had been studied before the onset of climate change, i.e. for *v* = 0. In this case *u*_inv_(*y*) = *ψ*(*y*) and, by Eq. (28), this function is for vulnerable species approximated by the observed distribution profile before environmental change. We denote the invasion fitness of the species for *v* = 0 by λ_0_. This quantity might also have been determined before the onset of environmental change, for example by measuring harvesting resistance (Rossberg 2013) or similar quantities. Assuming that the dispersal constant *D* is known as well, the fate of the population under environmental change (*v* ≠ 0) can now be predicted.

Specifically, since *ψ*(*y*) remains unchanged, invasion fitness will decline to:

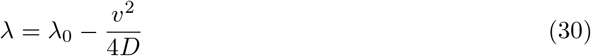

by Eq. (29). Remarkably, this decline is entirely independent of the form of the environmental performance curve. In particular, it does not depend on whether *g*(*y*) is left- or right-skewed. When environmental change is too fast 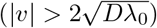, λ becomes negative and the species goes extinct.

For species that survive environmental change, the predicted population density is

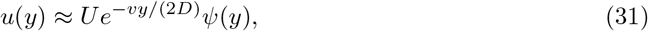

with

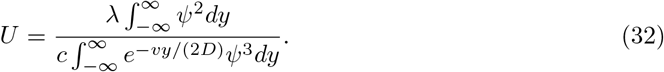

The factor *e*^−*vy*/(2*D*)^ in Eq. (31) shifts the value of *y* where *u*(*y*) is largest along the y axis, causing a lag between the actual distribution and the distribution one would expect in a static environment. The factor *e*^−*vy*/(2*D*)^ in the denominator in Eq. (32) corrects an artefact in Eq. (31) that arises when the maximum performance *g*(*y*) and so the maximum of *ψ*(*y*) are not close to *y* = 0. It has otherwise little effect.

### S1.5 Optimising the location of conservation interventions

Given the detrimental effect of environmental change, conservation ecologists have considered how a species’ population might best be supported to prevent extirpation during range shift. One management option is to support endangered populations by providing, e.g., additional food, shelter or nesting opportunities, suppressing competitors or natural enemies, or in case of exploited species, through targeted reductions in exploitation rates. A question that naturally arises is: where in a species range (between the leading edge and the trailing edge of the range) would such interventions be most effective? The first impulse might be, e.g., to support species in the leading edge of their shifting populations to accelerate the range shift. Interestingly, our theory suggests this isn’t always the best choice.

For simplicity, we assume that the conservation measures are introduced at a particular point *y*_cons_ along the *y*-axis (i.e the reference frame that moves with the climate) and reasonably represented in our model by modifying the environmental performance function *g*(*y*) to *g*(*y*) + *g*_cons_*δ*(*y* – *y*_cons_). Here a constant *g*_cons_ > 0 quantifies the strength (and sign) of the intervention or perturbation and *δ*(*y* – *y*_cons_) denotes Dirac’s delta-functional, a generalised function that represents a sharp peak that is localised to be non-zero only at *y* = ycons and is considered to have a ‘height’ such that ∫ *δ*(*y* – *y*_cons_)*dy* = 1.

We can evaluate the effect of this intervention perturbatively along the lines of the perturbation scheme above (S1.3). For this, we multiply the added term *g*_cons_*δ*(*y* – *y*_cons_)*u*(*y*) with the book-keeping parameter *ϵ* and include it amongst the “small” terms on the right in Eq. (20). The calculation then progresses as before. It leads to a modified result for the population scaling factor

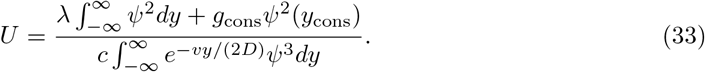

Thus, the effective invasion fitness resulting from these conservation measures increases to

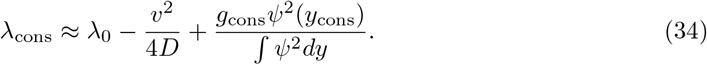

Hence, the conservation measures may indeed prevent extirpation of a population. Crucially, they are, by Eq. (34), most effective neither in the leading nor the trailing edge of the migrating population, but at the point where *ψ*(*y*) is largest. The mathematics underlying this result is closely related to that underlying the well-known sensitivity analyses for matrix models of age- or stage-structured population models (Caswell 2019). In the latter case, sensitivity is highest for matrix elements (*i, j*) for which the product of the reproductive value of stage *i* and numerical population size of stage *j* is largest. In our setting, the corresponding product evaluates to *ψ*^2^(*y*_cons_). This explains why it is neither efficient to support range-shifting populations at their leading edges nor at their trailing edges: the reproductive value of individuals at both ends is small and the density of surviving individuals small as well, which is why conservation measures aimed at the edges have little impact on the fate of the species as a whole. Remarkably, this rule is independent of the rate or direction of climate change and independent of the actual distribution of the migrating population.

### S1.6 Constraints on parameters for the Morse EPC model

Similar to the Schrödinger Equation (7), our equation for *ψ*(*y*) does not have ecologically valid solution for all parameter combinations. Here we discuss these limits on parameters in terms of the linear growth rate for *v* = 0, as given by Equation (10),

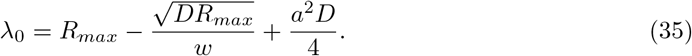

This expression attains a minimum in terms of *D* at *D* = 4*R_max_a*^−4^*w*^−2^. At this point λ = *R_max_*(1 – *a*^−2^*w*^−2^). This value of λ_0_ is the one that *g_Mor_*(*E*) approaches for large values of *aE*. It is negative by the condition |*aw*| < 1 that we imposed to assure that *g_Mor_*(*E*) declines to negative values for large |*E*|.

For *D* = 4*R_max_a*^−4^*w*^−2^, the solution *ψ*(*y*) of Eq. (3) does not decline to zero for either large positive or large negative *y*; rather, it describes the temporal decline of a population over a wide range in *y*. In the quantum physical analogue, this corresponds to a situations where, due to the quantum mechanical uncertainty principle, a potential well becomes unable to bind a quantum particles when it is too narrow and shallow. For values of *D* beyond this point Eq. (3) has no bounded solutions and hence cannot describe a growth from small abundance. In our analyses, we therefore always assume

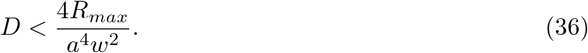

This constraint on *D* is a model artefact linked to the fact that *g_M or_* (*E*) approaches a constant value for large *aE*. Ecologically more realistic would be EPCs *g*(*E*) that always exhibit a steady decline with increasing |*E*| for both large positive and large negative *E*, in which case this constraint does not arise. For values of *D* well below the bound given by Eq. (36), however, this artefact of *g_M or_*(*E*) plays no role.

## S2 Mathematica Supplement: Solving Schrödinger equations for particular cases

The remainder of the mathematical results that stem from the use of the Schrödinger equation we present as an annotated and evaluated Mathematica notebook .pdf. Expressions used in the main text are highlighted with a light blue background. It is also available as `Mathematica.nb` in the code repository. While Mathematica is licensed software, this document can also ‘read’ using the free Wolfram player (https://www.wolfram.com/player/).

## S3 Model Specification

Separate .pdf file

## S4 Quantitative Match

Separate .pdf file

## References

Alexander, J.M., Chalmandrier, L., Lenoir, J., Burgess, T.I., Essl, F., Haider, S., et al. (2018). Lags in the response of mountain plant communities to climate change. Glob. Change Biol., 24, 563–579.

Berestycki, H., Diekmann, O., Nagelkerke, C.J. & Zegeling, P.A. (2009). Can a species keep pace with a shifting climate? Bull. Math. Biol., 71, 399–429.

Brito-Morales, I., García Molinos, J., Schoeman, D.S., Burrows, M.T., Poloczanska, E.S., Brown, C.J., et al. (2018). Climate Velocity Can Inform Conservation in a Warming World. Trends Ecol. Evol., 33, 441–457.

Brooker, R.W., Travis, J.M.J., Clark, E.J. & Dytham, C. (2007). Modelling species’ range shifts in a changing climate: The impacts of biotic interactions, dispersal distance and the rate of climate change. J. Theor. Biol., 245, 59–65.

Bull, J.W. (2015). Quantum Conservation Biology: A New Ecological Tool. Conserv. Lett., 8, 227–229.

Cantrell, R.S. & Cosner, C. (2003). Spatial Ecology via Reaction-Diffusion Equations. Wiley series in mathematical and computational biology. John Wiley & Sons Ltd.

Caswell, H. (2012). Matrix models and sensitivity analysis of populations classified by age and stage: a vec-permutation matrix approach. Theor. Ecol., 5, 403–417.

Caswell, H. (2019). Introduction: Sensitivity Analysis – What and Why? In: Sensitivity Analysis: Matrix Methods in Demography and Ecology, Demographic Research Monographs (ed. Caswell, H.). Springer International Publishing, Cham, pp. 3–12.

Chevin, L.-M., Lande, R. & Mace, G.M. (2010). Adaptation, Plasticity, and Extinction in a Changing Environment: Towards a Predictive Theory. PLoS Biol, 8, e1000357.

Elith, J. & Leathwick, J.R. (2009). Species distribution models: Ecological explanation and prediction across space and time. Annu. Rev. Ecol. Evol. Syst., 40, 677–697.

Fisher, R.A. (1937). THE WAVE OF ADVANCE OF ADVANTAGEOUS GENES. Ann. Eugen., 7, 355:369.

Grainger, T.N., Levine, J.M. & Gilbert, B. (2019). The Invasion Criterion: A Common Currency for Ecological Research. Trends Ecol. Evol., 34, 925–935.

Harsch, M.A., Phillips, A., Zhou, Y., Leung, M.R., Rinnan, D.S. & Kot, M. (2017). Moving forward: insights and applications of moving-habitat models for climate change ecology. J. Ecol., 105, 1169–1181.

Hastings, A. (2016). Timescales and the management of ecological systems. Proc. Natl. Acad. Sci., 113, 14568–14573.

Hastings, A., Cuddington, K., Davies, K.F., Dugaw, C.J., Elmendorf, S., Freestone, A., et al. (2005). The spatial spread of invasions: New developments in theory and evidence. Ecol. Lett., 8, 91–101.

Hoffmann, A.A. & Sgrò, C.M. (2011). Climate change and evolutionary adaptation. Nature, 470, 479–485.

Hurford, A., Cobbold, C.A. & Molnár, P.K. (2019). Skewed temperature dependence affects range and abundance in a warming world. Proc. R. Soc. B Biol. Sci., 286, 20191157.

Kareiva, P. & Shigesada, N. (1983). Analyzing insect movement as a correlated random walk. Oecologia, 56, 234–238.

Kerr, J.T. (2020). Racing against change: Understanding dispersal and persistence to improve species’ conservation prospects. Proc. R. Soc. B Biol. Sci., 287, 20202061.

Kot, M. & Phillips, A. (2015). Bounds for the critical speed of climate-driven moving-habitat models. Math. Biosci., 262, 65–72.

Legault, G., Bitters, M.E., Hastings, A. & Melbourne, B.A. (2020). Interspecific competition slows range expansion and shapes range boundaries. Proc. Natl. Acad. Sci., 117, 26854–26860.

Lenoir, J., Bertrand, R., Comte, L., Bourgeaud, L., Hattab, T., Murienne, J., et al. (2020). Species better track climate warming in the oceans than on land. Nat. Ecol. Evol., 4, 1044–1059.

Leroux, S.J., Larrivée, M., Boucher-Lalonde, V., Hurford, A., Zuloaga, J., Kerr, J.T., et al. (2013). Mechanistic models for the spatial spread of species under climate change. Ecol. Appl., 23, 815–828.

Levins, R. (1966). The strategy of model building in population biology. Am. Nat., 54, 421–431.

Levins, R. (1969). Some Demographic and Genetic Consequences of Environmental Heterogeneity for Biological Control1. Bull. Entomol. Soc. Am., 15, 237–240.

Li, B., Bewick, S., Shang, J.I.N. & Fagan, W.F. (2014). PERSISTENCE AND SPREAD OF A SPECIES WITH A SHIFTING HABITAT EDGE. SIAM J. Appl. Math., 74, 1397–1417.

Liu, B.R. & Kot, M. (2019). Accelerating invasions and the asymptotics of fat-tailed dispersal. J. Theor. Biol., 471, 22–41.

Lurgi, M., Brook, B.W., Saltré, F. & Fordham, D.A. (2015). Modelling range dynamics under global change: Which framework and why? Methods Ecol. Evol., 6, 247–256.

MacLean, S.A. & Beissinger, S.R. (2017). Species’ traits as predictors of range shifts under contemporary climate change: A review and meta-analysis. Glob. Change Biol., 23, 4094–4105.

Mattis, D.C. (1993). The Many-Body Problem: An Encyclopedia of Exactly Solved Models in One Dimension(3rd Printing with Revisions and Corrections). WORLD SCIENTIFIC.

Melbourne, B.A. & Hastings, A. (2009). Highly Variable Spread Rates in Replicated Biological Invasions: Fundamental Limits to Predictability. Science, 325, 1536–1539.

Morse, P.M. (1929). Diatomic Molecules According to the Wave Mechanics. II. Vibrational Levels. Phys. Rev., 34, 57–64.

Nadeau, C.P., Urban, M.C. & Bridle, J.R. (2017). Climates Past, Present, and Yet-to-Come Shape Climate Change Vulnerabilities. Trends Ecol. Evol., 32, 786–800.

Nagasawa, M. (1993). Equivalence of Diffusion and Schrödinger Equations. In: Schrödinger Equations and Diffusion Theory, Monographs in Mathematics (ed. Nagasawa, M.). Birkhäuser, Basel, pp. 89–114.

Parmesan, C. & Yohe, G. (2003). A globally coherent fingerprint of climate change. Nature, 421, 37–42.

Potapov, A.B. & Lewis, M.A. (2004). Climate and competition: The effect of moving range boundaries on habitat invasibility. Bull. Math. Biol., 66, 975–1008.

Real, R., Márcia Barbosa, A. & Bull, J.W. (2017). Species distributions, quantum theory, and the enhancement of biodiversity measures. Syst. Biol., 66, 453–462.

Ruel, J.J. & Ayres, M.P. (1999). Jensen’s inequality predicts effects of environmental variation. Trends Ecol. Evol., 14, 361–366.

Rumpf, S.B., Hülber, K., Wessely, J., Willner, W., Moser, D., Gattringer, A., et al. (2019). Extinction debts and colonization credits of non-forest plants in the European Alps. Nat. Commun., 10, 4293.

Savage, V.M., Gillooly, J.F., Brown, J.H., West, G.B. & Charnov, E.L. (2004). Effects of body size and temperature on population growth. Am. Nat., 163, 429–441.

Schrödinger, E. (1926). An Undulatory Theory of the Mechanics of Atoms and Molecules. Phys. Rev., 28, 1049–1070.

Skellam, A.J.G. (1951). Random Dispersal in Theoretical Populations. Biometrika, 38, 196–218.

Svenning, J.C. & Sandel, B. (2013). Disequilibrium vegetation dynamics under future climate change. Am. J. Bot., 100, 1266–1286.

Terry, J.C.D., O’Sullivan, J.D. & Rossberg, A.G. (2022). Synthesising the multiple impacts of climatic variability on community responses to climate change. Ecography, 2022, e06123.

Thompson, P.L. & Fronhofer, E.A. (2019). The conflict between adaptation and dispersal for maintaining biodiversity in changing environments. Proc. Natl. Acad. Sci. U. S. A., 116, 21061–21067.

Thompson, P.L. & Gonzalez, A. (2017). Dispersal governs the reorganization of ecological networks under environmental change. Nat. Ecol. Evol., 1, 0162.

Urban, M.C., Bocedi, G., Hendry, A.P., Mihoub, J.B., Peer, G., Singer, A., et al. (2016). Improving the forecast for biodiversity under climate change. Science, 353, aad8466.

Urban, M.C., Tewksbury, J.J. & Sheldon, K.S. (2012). On a collision course: competition and dispersal differences create no-analogue communities and cause extinctions during climate change. Proc. R. Soc. B Biol. Sci., 279, 2072–2080.

Yalcin, S. & Leroux, S.J. (2017). Diversity and suitability of existing methods and metrics for quantifying species range shifts. Glob. Ecol. Biogeogr., 26, 609–624.

Zhou, Y. & Kot, M. (2011). Discrete-time growth-dispersal models with shifting species ranges. Theor. Ecol., 4, 13–25.

Zurell, D., Jeltsch, F., Dormann, C.F. & Schröder, B. (2009). Static species distribution models in dynamically changing systems: How good can predictions really be? Ecography, 32, 733–744.

Zurell, D., Pollock, L.J. & Thuiller, W. (2018). Do joint species distribution models reliably detect interspecific interactions from co-occurrence data in homogenous environments? Ecography, 41, 1812–1819.

## Additional References

Al-Gwaiz, M.A. (2008). Sturm-Liouville Theory and its Applications. Springer Undergraduate Mathematics Series. Springer-Verlag, London.

Chen, L.-Y., Goldenfeld, N. & Oono, Y. (1996). Renormalization group and singular perturbations: Multiple scales, boundary layers, and reductive perturbation theory. Phys. Rev. E, 54, 376–394.

Rossberg, A.G. (2013). Food Webs and Biodiversity: Foundations, Models, Data. John Wiley & Sons Ltd, Oxford.

Stephenson, L.E. & Wollkind, D.J. (1995). Weakly nonlinear stability analyses of one-dimensional Turing pattern formation in activator-inhibitor/immobilizer model systems. J. Math. Biol., 33.

Zeidler, E. (1995). Applied Functional Analysis. Applied Mathematical Sciences. Springer New York, New York, NY.

