## Supplementary material for "Schrödinger’s range-shifting cat: analytic predictions for the effect of asymmetric environmental performance on climate change responses": SI 2

### Appendix 2

This document printed 30 June 2022. Made using Mathematica v.13.0

#### 2.1 Starting Assumptions

In[ ]:=
```
$Assumptions = (
    disp > 0 &&   (*Dispersal rate*)
      w > 0 &&     (*EPC width*)
      r0 > 0 &&    (*Maximum growth rate (at optima)*)
      v > 0 &&
      (*Speed of climate change --- we fix the sign of v  but keep that of a  open*)
      Element[y, Reals] &&    (*Spatial dimension, moving reference frame*)
      Element[a, Reals] &&    (*Asymetry of Morse EPC*)
      frontWidth > 0 &&    (* Property of population density shape,
      used to work around some Sqrt[] expressions. *)
      sigma > 0    (*standard deviation of high-frequency environmental variation*)
  )
```

Out[ ]= disp > 0 && w > 0 && r0 > 0 && v > 0 && y ∈ ℝ && a ∈ ℝ && frontWidth > 0 && sigma > 0

In[ ]:= (* Various rules used to simplify expressions later *)

dispRule = Solve$\left[\text{frontWidth} == \frac{\sqrt{\text{disp}}}{\sqrt{\text{r0}}}, \text{disp}\right]$ // Last

frontWidthRule = Solve$\left[\text{frontWidth} == \frac{\sqrt{\text{disp}}}{\sqrt{\text{r0}}}, \text{frontWidth}\right]$ // Last

r0Rule = Solve$\left[\text{frontWidth} == \frac{\sqrt{\text{disp}}}{\sqrt{\text{r0}}}, \text{r0}\right]$ // Last

Out[ ]= $\{\text{disp} \to \text{frontWidth}^2\, \text{r0}\}$

Out[ ]= $\left\{\text{frontWidth} \to \sqrt{\frac{\text{disp}}{\text{r0}}}\right\}$

Out[ ]= $\left\{\text{r0} \to \frac{\text{disp}}{\text{frontWidth}^2}\right\}$

#### 2.2 Defining Core Functions

Start by writing down the basic equation, without the non-linearity, in the co-moving reference frame. Here g() is the EPC,  f() is population density (called u_inv() in the paper), and  lam is the overall growth rate ($\lambda$).

Printed by Wolfram Mathematica Student Edition

Start by writing down the basic equation, without the non-linearity, in the co-moving reference frame. Here g() is the EPC, f() is population density (called u_inv() in the paper), and lam is the overall growth rate (λ).

In[•]:=
```
eq = lam*f[y] == g[y]×f[y] + v f'[y] + disp f''[y];
eq // TraditionalForm
```

Out[•]//TraditionalForm=

$$\text{lam}\, f(y) = \text{disp}\, f''(y) + v\, f'(y) + f(y)\, g(y)$$

#### 2.2.1 Morse (Asymmetric) EPC

In[•]:=
```
(* Define Morse Potential Function *)
gMorseRule = {g → Function[y, r0 (1 - 1/(a^2 w^2)×(1 - Exp[a y])^2)]}
(* Solve differential equation for f in terms of y.
   Rather than first computing ψ() and from this f() (u_inv() in the paper),
we here compute f() directly and let Mathematica to all the housekeeping.
   NB: The solution is very complex, so not showing here! *)
morseSol = DSolve[eq /. gMorseRule, f, y] // Last;
```

Out[•]=

$$\left\{g \to \text{Function}\left[y, r0\left(1 - \frac{(1 - \text{Exp}[a\, y])^2}{a^2\, w^2}\right)\right]\right\}$$

In[•]:=
```
(* Add additional condition for localisation of climatic niche
  (otherwise don't get positive growth anywhere and it blows up)*)
```

In[•]:=
```
(* Negativity of g()in the tails leads to additional conditions,
but we don't make them explicit in order not to confuse Mathematica.*)
Reduce[$Assumptions && 1/(a^2 w^2) > 1, a]
```

Out[•]=

$$y \in \mathbb{R}\ \&\&\ w > 0\ \&\&\ v > 0\ \&\&\ \text{sigma} > 0\ \&\&\ r0 > 0\ \&\&\ \text{frontWidth} > 0\ \&\&\ \text{disp} > 0\ \&\&\ \left(-\frac{1}{w} < a < 0\ \|\ 0 < a < \frac{1}{w}\right)$$

In[•]:=
```
(* The the solution we are looking for is that containing the
 Laguerre polynomial of zeroth order. Hence, we extract the order of the
 Laguerre polynomial and isolate the condition of it to be zero. *)
morseCond = 0 == (Cases[f[y] /. morseSol, LaguerreL[n_, _, _] → n, Infinity, 1] // First)
```

Out[•]=

$$0 == \frac{1}{2\, a^2\, \text{disp}\, w}\left(2\sqrt{\text{disp}\, r0} - a^2\, \text{disp}\, w - \sqrt{\left(4\, \text{disp}\, r0 + 4\, a^2\, \text{disp}\, \text{lam}\, w^2 - 4\, a^2\, \text{disp}\, r0\, w^2 + a^2\, v^2\, w^2\right)}\right)$$

##### 2.2.2 Harmonic EPC

In[ ]:=
```
(*Also consider the harmonic EPC. Needed below for baseline calculations of shift*)
(*Follows same logic.*)
gHarmRule = {g → Function [y, r0 (1 - y^2 / (w^2))]} (* Harmonic Potential *)
harmSol = DSolve[eq /. gHarmRule, f, y] // Last;
harmCond = 0 == (Cases[f[y] /. harmSol, HermiteH[n_, _] → n, Infinity, 1] // First);
harmLamRule = Solve[harmCond, lam] // Last // FullSimplify;
(* Now compute the solution for the case
 where the Hermite polynomical is of order zero.*)
harmSolClean = {f → Function @@
     {y, f[y] /. harmSol /. harmLamRule /. dispRule /. c1 → 1 /. c2 → 0 // FullSimplify}};
```

Out[ ]= $\left\{g \to \text{Function}\left[y,\ r0\left(1-\frac{y^2}{w^2}\right)\right]\right\}$

#### 2.3 Response 1: Population Invasion Fitness & Critical Speed of Climate Change

In[ ]:=
```
(* Get lambda from morseCond. *)
morseLamRule = Solve[morseCond, lam] // Last // FullSimplify ;
 Style[%, Background -> LightBlue]
```

Out[ ]= $\left\{\text{lam} \to \frac{a^2\,\text{disp}}{4} + r0 - \frac{v^2}{4\,\text{disp}} - \frac{\sqrt{\text{disp}\ r0}}{w}\ \ \text{if disp} < \frac{4\,r0}{a^4\,w^2}\right\}$

In[ ]:=
```
(* Create a neater solution, specifically for the case given by morseCond,
for future use. Assume that the condition above is satisfied.*)
morseSolClean = {f → Function @@ {y,
      Assuming[-2 + a^2 frontWidth w < 0,

        f[y] /. morseSol /. morseLamRule /. dispRule /. c1 → 0 /. c2 → 1 // FullSimplify]
     }}
```

Out[ ]= $\left\{f \to \text{Function}\left[y,\ e^{-\frac{\left(2\,e^{a\,y}\,\text{frontWidth}\ r0+a\left(-2\,\text{frontWidth}\ r0+a\left(a\,\text{frontWidth}^2\ r0+v\right)w\right)\right)y}{2\,a^2\,\text{frontWidth}^2\ r0\ w}}\right]\right\}$

##### 2.3.1 Critical Speed of Climate Change

In[ ]:=
```
(* All we need here is lambda_0, the rest is trivial *)
lam0Rule = (lam0 -> lam) /. morseLamRule /. v -> 0 // FullSimplify;
Style[lam0Rule, Background -> LightBlue]
```

Out[ ]= $\text{lam0} \to \frac{a^2\,\text{disp}}{4} + r0 - \frac{\sqrt{\text{disp}\ r0}}{w}\ \ \text{if disp} < \frac{4\,r0}{a^4\,w^2}$

#### 2.3.2 Impact of Weather (high frequency variation in growth rate)

In[ ]:=
```
(* Convolution of Morse performance curve with Gaussian weather variability gives another,
modified Morse performance curve: *)
gMorseWeather =
  Integrate[(g[x] /. gMorseRule) * PDF[NormalDistribution[0, sigma]][y - x],
   {x, -Infinity, Infinity}, GenerateConditions -> False]
```

Out[ ]=

$$\frac{r0\left(-1-e^{2 a\left(a\, sigma^2+y\right)}+2\, e^{\frac{a^2 sigma^2}{2}+a y}+a^2 w^2\right)}{a^2 w^2}$$

#### 2.3.3 How effective values of key parameters are impacted by weather

In[ ]:=
```
(* Because doing it directly takes too long, so we break it up into a series expansion *)
gWSol1 =
    Solve[And @@ Table[SeriesCoefficient[
                Series[(g[y] /. gMorseRule /. {r0 -> r02, w -> w2, y -> y - y2}) - gMorseWeather,
                        {y, 0, 2}],
                    n] == 0,
            {n, 1, 1}],
        {r02}] // FullSimplify // Last;


gWSol2 = Assuming[Element[y2, Reals],
            Solve[And @@ Table[SeriesCoefficient[Series[
                (g[y] /. gMorseRule /. {r0 -> r02, w -> w2, y -> y - y2}) - gMorseWeather,
                        {y, 0, 2}],
                    n] == 0,
                {n, 2, 2}] /. gWSol1, {y2}]] // Last;


gWSol3 = Assuming[Element[y2, Reals] && w2 > 0 && a > 0,
        Solve[And @@ Table[SeriesCoefficient[Series[(g[y] /. gMorseRule /.
                    {r0 -> r02, w -> w2, y -> y - y2}) - gMorseWeather,
                        {y, 0, 2}],
                    n] == 0,
                {n, 0, 0}] /. gWSol1 /. gWSol2 /. w2 -> Sqrt[w4] // FullSimplify,
        {w4}]] // Last // FullSimplify;
```

In[ ]:=
```
morseWeatherRule =
 Assuming[a > 0 (* This is just not to confuse Mathematica *), {r0 → r02,
     frontWidth → Sqrt[r0 / r02] * frontWidth,
     y → y - y2,
     w → Abs[w2]
    } /. gWSol1 /. gWSol2 /. w2 → Sqrt[w4] /. gWSol3 // FullSimplify]
```

Out[ ]= $\left\{r0 \to \frac{r0\left(-1+e^{-a^2\,sigma^2}+a^2\,w^2\right)}{a^2\,w^2},\ frontWidth \to a\,e^{\frac{a^2\,sigma^2}{2}}\,frontWidth\,w\,\sqrt{\frac{1}{1+e^{a^2\,sigma^2}\left(-1+a^2\,w^2\right)}},\ y \to \frac{3\,a\,sigma^2}{2}+y,\ w \to \sqrt{Abs\left[\frac{1+e^{a^2\,sigma^2}\left(-1+a^2\,w^2\right)}{a^2}\right]}\right\}$

In[ ]:=
```
(* Check consistency of calculation: *)
Assuming[a > 0 (* This is just not to confuse Mathematica *),
  morseWeatherRule /. sigma → 0 // FullSimplify]
```

Out[ ]= {r0 → r0, frontWidth → frontWidth, y → y, w → w}

##### 2.3.4 How weather affects population fitness:

In[ ]:=
```
(* As an example, expand dependence of max local fitness up to 4th orderin sigma *)
Series[r0 /. morseWeatherRule, {sigma, 0, 4}]
```

Out[ ]= $r0-\frac{r0\,sigma^2}{w^2}+\frac{a^2\,r0\,sigma^4}{2\,w^2}+O[sigma]^5$

In[ ]:=
```
(*Use above results to compute a new overall expression
  for growth rate that includes the effect of weather (sigma) *)
(*For simplicity include variety of assumptions*)
morseSigWeatherRule =
  Assuming[a > 0, (Assuming[disp < 4 r0/(a^4 w^2), morseLamRule // FullSimplify] /. morseWeatherRule //
       FullSimplify)] /. Sign[1 + e^(a^2 sigma^2) (-1 + a^2 w^2)] → 1 // FullSimplify
```

Out[ ]= $\left\{lam \to \frac{a^2\,disp}{4}+r0-\frac{v^2}{4\,disp}-\sqrt{\frac{disp\,e^{-a^2\,sigma^2}\,r0}{w^2}+\frac{\left(-1+e^{-a^2\,sigma^2}\right)r0}{a^2\,w^2}}\right\}$

In[ ]:=
```
(* Does weather increase or decrease lambda? *)
weatherDirRule = D[(lam /. morseSigWeatherRule), {sigma, 2}] /. sigma → 0 // FullSimplify;
Style[weatherDirRule, Background -> LightBlue]
```

Out[ ]= $\frac{-2\,r0+a^2\,\sqrt{disp\,r0}\,w}{w^2}$

#### 2.4 Response 2 - Lags

### 2.4 Response 2 - Lags

We analyze two measures of lag behind climate velocity : 1) the location of the peak density, and 2) the center of mass of the population.

#### 2.4.1: Lag of peak

In[∘]:=
```
(*Solve for location of peak of population density in terms of y (the moving window) *)
(* Harmonic EPC, that's easy: *)
harmShift = y /. Solve[D[f[y] /. harmSolClean, y] == 0, y] // Last // FullSimplify


(*Morse EPC*)
(* Firsts, compute the position of peak under climate change,
disregarding that without climate changes is also not at zero. *)
morseShiftRaw = y /. Solve[D[f[y] /. morseSolClean, y] == 0, y] // Last // FullSimplify
(* Now, compute compute actual shift and remove conditions. *)
morseShift = morseShiftRaw - (morseShiftRaw /. v → 0) // FullSimplify // Normal
(* Take first derivtive in v at v==0,
which corresponds to the (negative of the) time lag when v is small. *)
Series[D[morseShift, v] /. v → 0, {a, 0, 2}] /. frontWidthRule // FullSimplify
```

Out[∘]= $-\frac{v\,w}{2\,\text{frontWidth}\,r0}$

Out[∘]= $\frac{\text{Log}\left[1-\frac{a\,(a\,\text{frontWidth}^2\,r0+v)\,w}{2\,\text{frontWidth}\,r0}\right]}{a}$ if *condition*

Out[∘]= $\frac{1}{a}\left(-\text{Log}[\text{frontWidth}\,r0]+\text{Log}\left[\frac{-2\,\text{frontWidth}\,r0+a\,(a\,\text{frontWidth}^2\,r0+v)\,w}{-2+a^2\,\text{frontWidth}\,w}\right]\right)$

Out[∘]= $-\frac{w}{2\sqrt{\text{disp}\,r0}}-\frac{w^2\,a^2}{4\,r0}+O[a]^3$

In[ ]:=
```
(* Effect of weather on "friction" constant: *)
Assuming[a > 0, Series[D[morseShift // FullSimplify, v] /. v → 0, {a, 0, 2}] // FullSimplify]
(* without 'weather'*)
Assuming[e^(a^2 sigma^2) (-1 + a^2 w^2) > -1, Series[D[morseShift /. morseWeatherRule // FullSimplify, v] /.
        v → 0, {a, 0, 2}] // FullSimplify] /. frontWidthRule //
    FullSimplify // Normal // PowerExpand // FullSimplify // Apart;
Style[%, Background -> LightBlue]
```

Out[ ]= $-\frac{w}{2\,(\text{frontWidth r0})} - \frac{w^2 a^2}{4\,\text{r0}} + O[a]^3$

Out[ ]= $\begin{cases} \frac{\sqrt{\text{disp r0}}\, w}{2\,\text{disp r0}} + \frac{a^2\sqrt{\text{disp r0}}\,\text{sigma}^2 w}{4\,\text{disp r0}} - \frac{a^2 w^2}{4\,\text{r0}} & a \le 0 \\ -\frac{(2+a^2\,\text{sigma}^2)\, w}{4\sqrt{\text{disp}}\sqrt{\text{r0}}} - \frac{a^2 w^2}{4\,\text{r0}} & \text{True} \end{cases}$

#### 2.4.2: Centre of mass lag:

In[ ]:=
```
(* First compute the total area under the curve, to standardise with *)
morseNorm = Integrate[f[x] /. morseSolClean,
      {x, -Infinity, Infinity}, GenerateConditions → False] // FullSimplify;


(* Compute the moment generating function (using t as dummy argument of the MGF). *)
morseMGF = Integrate[Exp[t y] f[y] /. morseSolClean, {y, -Infinity, Infinity},
        GenerateConditions → False] / morseNorm // FullSimplify;


(* Compute the first moment, i.e. the centre of mass. *)
morseCentreMass = D[morseMGF, t] /. t → 0 // FullSimplify;
```

The formula for the position of the centre of mass looks singular for v, a == 0, but actually it’s smooth (technically, it can be “analytically continued” to $a==0$ and $v==0$). Based on the symmetry of the problem, centre of mass (in the co-moving reference system) is given by \text{centre of mass} = c_1 a + c_2 v + c_3 a^3 + c_4 a^2 v + c_5 a v^2 + c_6 v^3 + terms of order 5 and higher, with some constants c_1, ..., c_6.
However, here we are only interested in the lag in time for small v. This can be computed as D[\text{centre of mass},v] (at v=0) = c2 + c4 a^2 + terms of order 4 and higher. Since $c2$ is the same as for the harmonic potential, we take if from there.
To get $c4$, we first compute (1/2) * d^2 (\text{centre of mass}) / da^2 = 3 c_3 a + c_4 v + terms of order 2 and higher (and calls this “morseCentreMassEffect”), then takes the limit a ->0 (which remove the c_3 term and all higher order terms containing a) and then takes the derivative with respect to v at v = 0 (which converts c_4 v to c_4 and drops the remaining higher order terms). The result is c_4.

Finally, the lag c_2 + c_4 a^2 is assembled from the isolated terms.

In[∘]:=

```
morseCentreMassEffect = (1 / 2) D[morseCentreMass, {a, 2}] // FullSimplify;
morseCentreMassCorrection =
  a ^ 2 D[Limit[morseCentreMassEffect, a → 0, Direction → "FromAbove"] // FullSimplify, v] /.
    v → 0
harmLag = D[y /. Solve[D[f[y] /. harmSolClean, y] == 0, y] // Last, v];

morseCentreMassLag = harmLag + morseCentreMassCorrection // FullSimplify;

(*Present in a neater form*)
morseCentreMassLag /. frontWidthRule // FullSimplify // Apart

(* Now include the effect of weather variation.
       With some corner case-avoiding assumptions,
 find the power series expansion in terms of a to second order,
and present using raw parameters. *)
morseCentreMassLag2 =
  Assuming[1 + ⅇ^(a^2 sigma^2) (-1 + a^2 w^2) > 0,
          (Series[morseCentreMassLag /. morseWeatherRule // FullSimplify, {a, 0, 2}] //
              FullSimplify)] /. frontWidthRule // PowerExpand // Normal // Apart
(* At this order, the results turns out to be identical to that for the lag of the peak. *)
```

Limit: Warning: Assumptions that involve the limit variable are ignored.

Out[∘]= $-\frac{a^2 w^2}{2\, r0}$

Out[∘]= $-\frac{\sqrt{disp\, r0}\; w}{2\, disp\, r0} - \frac{a^2 w^2}{2\, r0}$

Out[∘]= $\begin{cases} -\frac{(2+a^2\, sigma^2)\, w}{4\sqrt{disp}\,\sqrt{r0}} - \frac{a^2 w^2}{2\, r0} & a > 0 \\ \frac{(2+a^2\, sigma^2)\, w}{4\sqrt{disp}\,\sqrt{r0}} - \frac{a^2 w^2}{2\, r0} & True \end{cases}$

### 2.5 Response 3: Local of Maximum Sensitivity

In[ ]:=
```
(* Calculation of location (in terms of y) of maximum sensitivity  *)
sensitivity = (f[y] /. morseSolClean /. v → -v) * (f[y] /. morseSolClean) // FullSimplify;
ySenseRule = Solve[D[sensitivity, y] == 0, y] // FullSimplify // Last;

Style[ySenseRule, Background -> LightBlue]
```

Out[ ]= $\left\{y \to \frac{\text{Log}\left[1 - \frac{1}{2} a^2 \text{ frontWidth } w\right]}{a} \text{ if } w < \frac{2}{a^2 \text{ frontWidth}}\right\}$

In[ ]:=
```
(*Calculate f at y =0, to standardise plotting height*)
f0 = f[y] /. morseSolClean /. y → 0
```

Out[ ]= $e^{-\frac{1}{a^2 \text{ frontWidth } w}}$

In[ ]:=
```
Plot[{ g[y] /10 /. gMorseRule,  (* Plot out Growth function [Blue] *)
        f[y]/f0 /. morseSolClean,
      (* Plot out psi function, standardised to equal 1 at y=0 [Orange ]*)
      sensitivity/f0^2} /.        (* Plot out Sensitivity, also scaled [Green]*)
     {a → 1.4, w → 0.5, frontWidth → 0.9, v → 4, r0 → 6} // Evaluate,
  {y, -5, 5},  (* Define conditions to plot*)
     PlotRange → {-1, All}, (* vertical axis range *)
    PlotLabel -> "a = 1.4",
     PlotLegends -> Placed[LineLegend[ColorData[97, "ColorList"][[1 ;; 3]],
       {"EPC", "Density", "Sensitivity"}], {0.85, 0.8}]]

Plot[{ g[y] /10 /. gMorseRule,
        f[y]/f0 /. morseSolClean,
      sensitivity/f0^2} /.
     {a → -1.4, w → 0.5, frontWidth → 0.9, v → 4, r0 → 6} // Evaluate, {y, -5, 5},
     PlotRange → {-1, All},
    PlotLabel -> "a = -1.4",
     PlotLegends -> Placed[LineLegend[ColorData[97, "ColorList"][[1 ;; 3]],
       {"EPC", "Density", "Sensitivity"}], {0.85, 0.8}]]
```

Out[∘]=

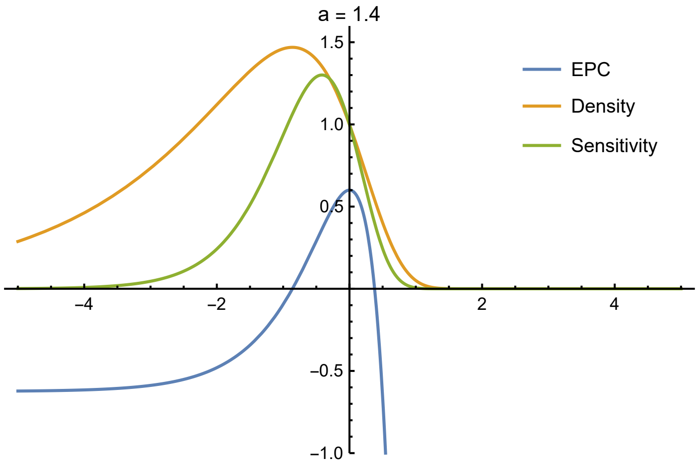


General: Exp[−1235.37] is too small to represent as a normalized machine number; precision may be lost.

General: Exp[−2474.85] is too small to represent as a normalized machine number; precision may be lost.

Out[∘]=

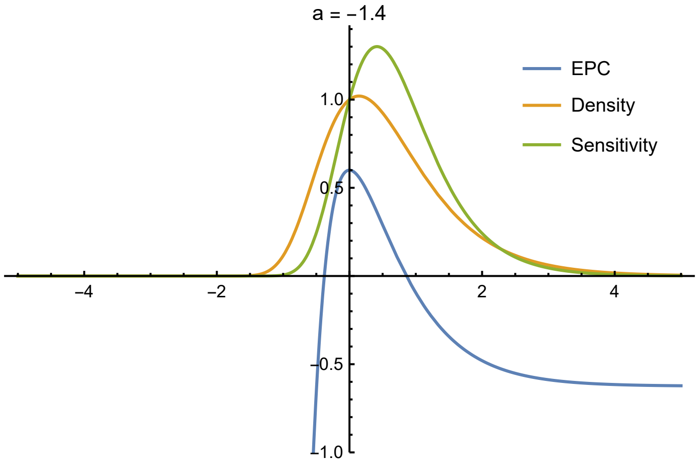


In[∘]:=
```
Plot[y /. ySenseRule /. frontWidth → 1 /. w → 1 // Evaluate, {a, −2, 2},
  AxesLabel -> {"Asymmetry", "Location of Peak Sensitivity"},
      PlotLabel -> "Location of Peak Sensitivity in terms of asymmetry\nPeak sensitivity
       depends on shape of curve, not on \n direction of climate change" ]
```

Out[∘]=

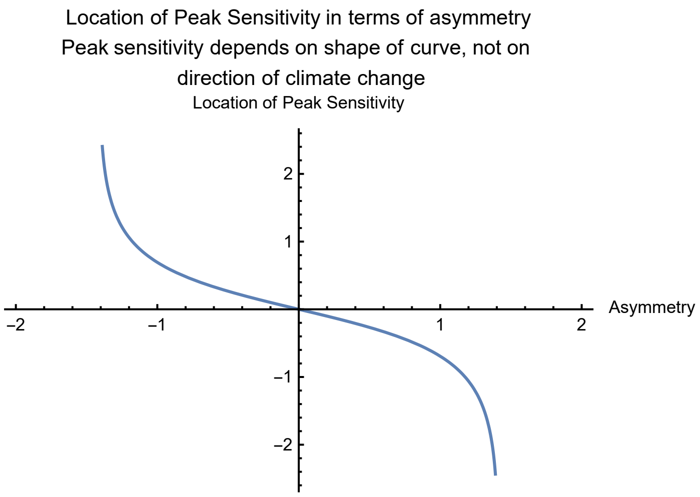


#### 2.6 Example Plots

### 2.6 Example Plots

In[ ]:=
```
num = Join[dispRule, {w → 1, r0 → 1, a → 3/4, v → 0.5, frontWidth → 0.6}];
morseLamRule //. num
(*Repeatedly put in the parameters to find what lambda is with these*)
Plot[g[y] /. gMorseRule //. num, {y, -7, 7},
      PlotRange → {-r0, r0} //. num // Evaluate,
      PlotLabel -> "Shape of EPC"]
Plot[f[y] / Abs[morseNorm] /. morseSolClean //. num // Evaluate, {y, -7, 7},
      PlotRange → All,
      PlotLabel -> "Shape of population density"]
```

Out[ ]= {lam → 0.277014}

Out[ ]=

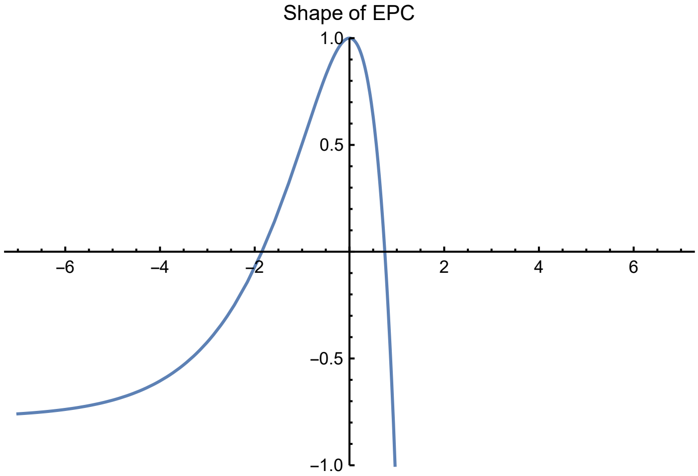


Out[ ]=

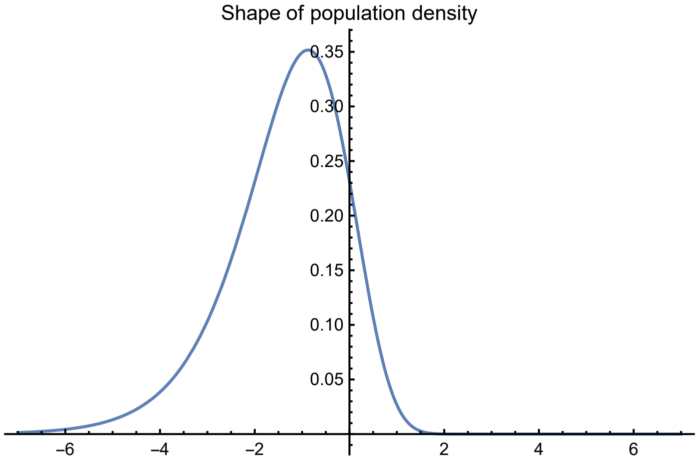


In[ ]:=
```
(* Evaluation through time*)
(* Specify movement equation explicitly,
then put into moving reference frame. Climate change starts at t==0. *)
```

In[ ]:=
```
eqPLOT = (D[f[x - x0[t], t], t] == disp * D[f[x - x0[t], t], {x, 2}] +
        g[x - x0[t]] × f[x - x0[t], t] - f[x - x0[t], t]^2) /. x → y + x0[t];
L = 13; (* 0.5 * spatial extent *)
tMin = -20;
tMax = 25;
num =
   Join[gMorseRule,
    {disp → 0.2,
        v0 → 0.45, (*rate of climate change*)
      a -> 0.9,
      w -> 1,
      r0 -> 1,    x0 → Function[t, t * v0 * HeavisideTheta[t]] }];

sol = NDSolve[eqPLOT &&
              f[y, tMin] == (11 * f[y] /. morseSolClean /. v -> 0 /. frontWidthRule) &&
         (*Set starting distribution to 0.2*(1-(1-Exp[y])^2)  *)
              f[-L, t] == 0 &&
              f[L, t] == 0 //. num // FullSimplify,
           f, (*Solve for f*)
    {y, -L, L}, (*Range of y to consider*)
    {t, tMin, tMax}(*Range of times to consider*)
   ] // Last
```

Out[ ]=
```
{f → InterpolatingFunction[ Domain: {{-13., 13.}, {-20., 25.}}
                            Output: scalar ]}
```

In[ ]:=
```
pl1 = ContourPlot[10^-2 + f[x, t] /. sol, {t, 0.2 tMin, tMax},
                  {x, -5, 2},
                 PlotRange → {0, All}, PlotPoints → 50, FrameLabel → {"Time", "y"}];

(*Add boundary for where population is above a threshold*)
pl1r = ContourPlot[10^-2 + f[x, t] /. sol,
                  {t, 0.2 tMin, tMax},
                  {x, -5, 2},
                  PlotRange → {0, All}, PlotPoints → 50, Contours → {0.1},
                  ContourStyle → {{Red, Thick}}, ContourShading → None];

(*Location of peak of distribution*)
pl3 = ListPlot[Table[{t, y /. FindMaximum[f[y, t] /. sol,
                         {y, -0.1 L, 0.1 L}][[2]]},
                    {t, 0.2 tMin, tMax, 0.1}], Joined → True];

(*Combine all the parts of the plot together*)
Show[pl1, pl3, pl1r]
```

Out[ ]=

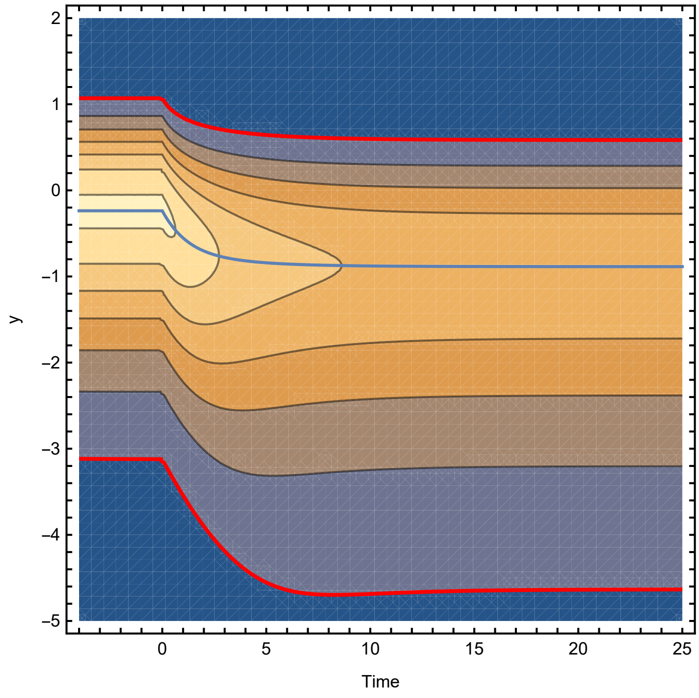

In[ ]:=
```
(* Same plot, now in spatially fixed reference frame *)
(* Define f for fixed coordinate system *)
fFixed[x_, t_] = If[Abs[x - x0[t]] < L, 10^-2 + f[x - x0[t], t], 0] //. num;

(* Contour plot of f *)
pl1 = ContourPlot[fFixed[x, t] /. sol // Evaluate, {t, 0.2 tMin, tMax},
    {x, -5, 2},
    PlotRange → {0, All}, PlotPoints → 50, FrameLabel → {"Time", "x"}];

(*Add boundary for where population is above a threshold*)
pl1r = ContourPlot[10^-2 + fFixed[x, t] /. sol // Evaluate,
    {t, 0.2 tMin, tMax},
    {x, -5, 2},
    PlotRange → {0, All}, PlotPoints → 50, Contours → {0.1},
    ContourStyle → {{Red, Thick}}, ContourShading → None];

(*Location of peak of distribution*)
pl3 = ListPlot[Table[{t, (x0[t] //. num) + y /. FindMaximum[f[y, t] /. sol,
        {y, -0.1 L, 0.1 L}][[2]]},
      {t, 0.2 tMin, tMax, 0.1}],
   Joined → True];

(*Combine all the parts of the plot together*)
Show[pl1, pl3, pl1r]
```

Out[∘]=

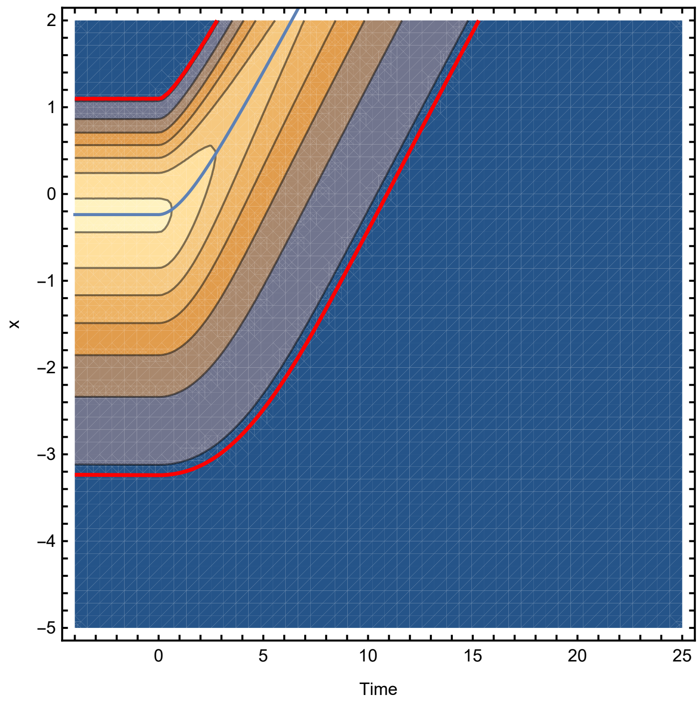
