## Supplementary material for "Schrödinger’s range-shifting cat: analytic predictions for the effect of asymmetric environmental performance on climate change responses": SI 3

### SI 3: Simulation Model Specification

#### Introduction

This appendix provides further background to the simulation model used to test the potential applicability of the analytic results to situations that include a more complex and heterogeneous spatial and temporal structure. All code is publicly available (as described in the data availability statement). References to scripts given in `typewriter font` are designed to point the interested reader to specific functions in cases where further detail is needed. `SimulationRun.rmd` conducts the assembly and tests, results are analysed in `Plots.rmd`.

#### Overall Setting

The model describes the population of each species at a set of specific sites (nodes) distributed in a rectangular arena. Each node is assigned a particular environmental value at each point in time. The model is advanced in discrete time. In this simulation the species do not interact, so strictly it is a set of metapopulations, rather than a genuine metacommunity. However, we will use the terms 'community' or 'assembly' to refer to the set of species present in a particular model run.

There are two important time scales. We refer to the slower timescale as 'years' and this is the timescale on which the environment changes. There is also a finer timescale (1/15th year), that is used to progress the population dynamics model (see below).

Where parameters are not hard coded into the model functions they are specified in `Parameters/Parameters.R`, and where relevant described in the text below.

#### Spatial Setting

Each assembly has a random spatial distribution of 200 nodes (See figure below for an example), whose $x$ and $y$ coordinates are drawn from a uniform distribution of from 0 to 40 and 0 to 10 respectively. This process generates arenas with notable spatial heterogeneity.

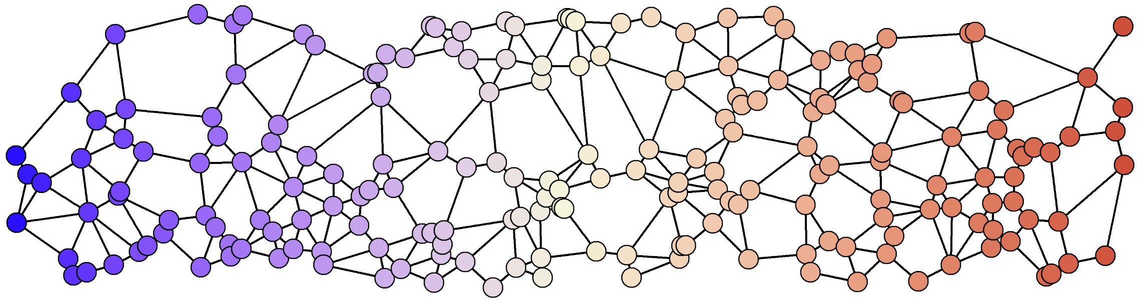

#### Environment and climate change

Each node is assigned an environmental variable, initially equal to its $x$ value. Each year this is subject to up to two forms of variation:

1. Stochastic variation. This is additive and drawn from a Gaussian distribution ($\sigma = 0.5$), and applied universally across the arena.
2. Climate change. After the date of the onset of climate change, the value of $E$ is increased by a climate change rate value $v$ at the start of each 'year'.

#### Population Growth rate

Each population has its own unique optimal environmental value ($\phi$), but all share the same environmental performance function based on the Morse potential function (`RCalc.R`). For species $i$ in site $x$ at time $t$:

$$R_{i,x,t} = R_{max}\left(1 - \frac{\left(1 - e^{a(E_{x,t}-\phi_i)}\right)^2}{a^2 w^2}\right)$$

with the additional constraint that $R_{i,x}$ is lower bounded at -100. Here, $R_{max}$ = 12, $a$ = -0.9 or 0.9, $w$ = 1.

#### Node Connectance

Nodes are determined to be connected with the Gabriel algorithm (see figure above), that determines the closest node in each direction (`genAdjMat()` in `NetworkGenerators.R`).

#### Dispersal between nodes

A matrix of between-node dispersal rates is generated from the distance between nodes and some key parameters: migration rate $m$, dispersal length $L$, and normalised so that the total emigration is constant.

If the Euclidean distance between two connected nodes A and B = $d_{a,b}$, then the dispersal rate of B to A was

$$D_{a,b} = m\frac{e^{-d/L}}{D^*},$$

where the normalising term $D*$ sums up all the outgoing migration from site B to the set of nodes $n$ that it is connected to

$$D^* = \sum_n e^{-d_{n,b}/L}$$

The dispersal matrix also includes negative diagonal terms representing emigration:

$$D_{a,a} = -m.$$

#### Population dynamics within each node

The core population dynamics within each node is a very simple discrete time model (`metacDynamics.R`):

For a number of time steps ($t_{max}$), the population ($B$) at that node ($z$) in the next step is

$$B_{z,t+1} = B_{z,t} + B_{z,t}(R - B_{z,t}) + \sum_n D_{z,n} B_{n,t},$$

where $n$ runs over the set of all nodes, although in practice only the neighbours contribute.

To avoid negatives and ongoing numerical errors, at each step any populations that were to fall below 0 are set to 0.

In this simulation, 15 of these small time steps $t$ are taken each 'year', all under the same environmental conditions.

### Community Asssembly and 'burn-in'

#### Generation of new species

Each assembly is initiated by generating 100 species (`PopulateSystem.R`). Each is assigned an environmental optimum ($\phi$) based on a random uniform distribution bounded from 20 to 30. Each species is then introduced at density 10 at the node closest to this optimum. A burn in period of 200 years is then run, in which the species can expand to fill their range.

#### Removal of extinct species

At the end of each 'year', any species that are not above a specified extinction threshold ($10^{-6}$) in at least one node are removed from the simulation (`clean_extinct.r`).

### Tests

Once the communities are assembled (one set of communities with 'warm-skewed' EPCs and the other with 'cold-skewed' EPCs, based on the sign of $a$), two independent tests of the effect of climate change are conducted: 1) the lag between the climate shift and the population shift, 2) the speed of climate change that the species cannot cope with (critical rate).

#### Lags

The lags test is run using the function in `LagMeasurer.R`. The core response is the weighted mean x-coordinate of the population distribution across the nodes, the mean population location.
To determine the starting location, the dynamics are run for 20 'years' with underlying stochastic climate variation but without climate change, and an average taken.

Then climate change is introduced at a rate of 0.1 units per year, and 50 years of simulation run to overcome any transient effects. During this period the removal of extinct species is switched off - in practice this makes little difference, but prevents occasional extinctions disrupting the tracking of species movements. The climate change is continued, and the mean population location recorded over 20 subsequent years. Lastly, the total amount of climate change at each of the recording years is compared to the observed population displacement and averaged. Recall that, since the initial environment is given by the $x$-spatial coordinate, there is a direct correspondence. The total of 7 units ($0.1 \times 50+20$ ) of climate change would not be enough to push any species off the edge of the arena.

A linear mixed effects model was used to estimate the effect of the direction of EPC asymmetry on the species lag. Assemblage ID was included as a random effect as the species within each assemblage shared a stochasticity pattern and a spatial network. The total number of observations was 19948, across 200 assemblages. The fitted assemblage random effect explained a variance of 0.03036, with 0.02912 residual variance remaining. The overall fitted intercept value was 0.46736 (SE= 0.01751 ) and the fixed effect of $a = 0.9$ was 0.15067 (SE = 0.02476).

#### Critical Speed of Climate Change

The critical climate change rate tests was conducted with `CritCC_Calculator.R`. The core response is the proportion of species that go extinct at series of rates of climate change. We tested 50 'years' of climate change at rates of 0 to 0.4, in steps of 0.025. Thresholding was removed, instead we looked at the proportion of species within each assemblage that fell below 1.0 across all nodes (other measures of 'extinction' gave similar results).

The impact of EPC skew direction was estimated using a logistic generalised linear mixed effects model. The rate of climate change was also included as a main effect including an interaction term. The number of species in each assemblage was used as a weighting term and the assemblage ID was incorporated as a random effect. In R syntax this was: `lme4::glmer(Data,  FracExtant1 ~  CC_rate*factor(EPC_A) + (1|AssemblageID), family = 'binomial',  weights = NumberFocalStart )`

The full model results were:

| | Estimate | Std. Error | z value | p-value |
|---|---|---|---|---|
| Intercept | 22.5680 | 0.2484 | 90.85 | <2e-16 *** |
| CC_rate | -74.0872 | 0.5023 | -147.48 | <2e-16 *** |
| EPC a =0.9 | -4.2961 | 0.3563 | -12.06 | <2e-16 *** |
| CC_rate:EPC =0.9 | 11.4094 | 0.7107 | 16.05 | <2e-16 *** |

*Table S3.1* Model fits for the critical rate of climate change.

The assemblage ID random effect (over 200 assemblages) had a SD of 2.088 and the total the number of trials was 2600.

#### Sensitivity to Conservation Intervention.

The trials of different intervention locations were conducted with `Peak_Sensitivity_Calc()`. The same assemblages were used and climate change was introduced at a rate of $v = 0.28$, a rate chosen based on the results of the critical rate of climate change trials described above.

The 'conservation intervention' was modeled as a +2 increase of the underlying population growth rate ($R_{i,x,t}$) at particular sites. Sites were initially modified when they had an x-coordinate that fell within the conservation region that was determined for each species based on the location of the optimum, and an offset parameter $L$. As climate change progressed, this window of conservation intervention moved with the climate velocity. Examples of growth curves modified in this way with different skew directions ($a$) and locations of interventions ($L$) are given in the figure below:

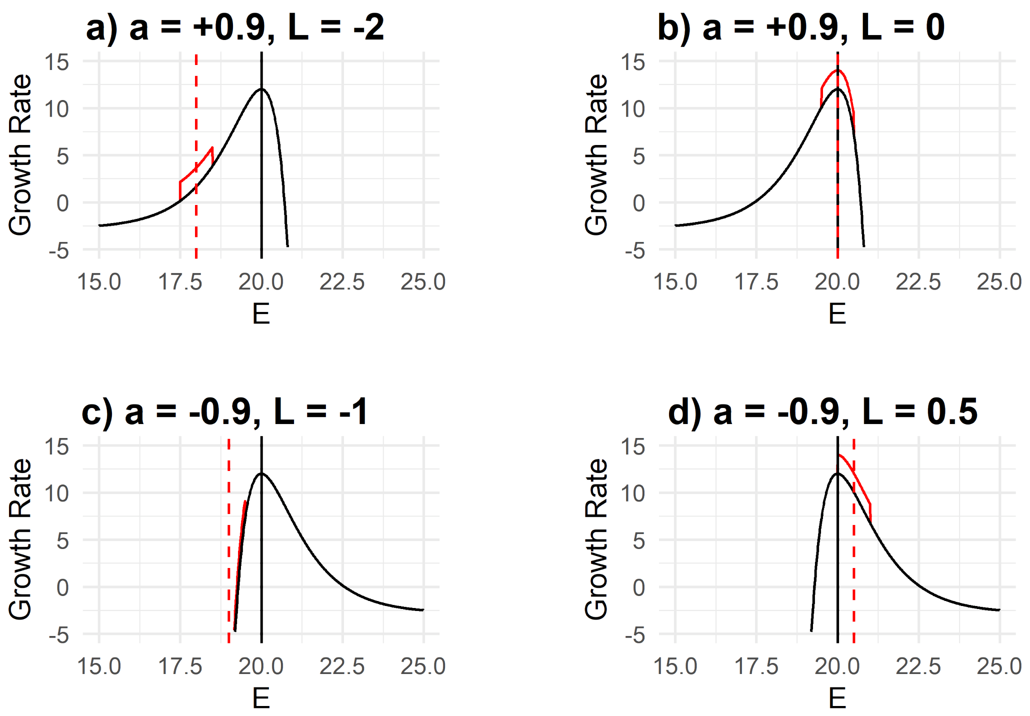


**Figure S3.1** Illustration of how conservation impacts were modelled by increasing the growth rates in an intervention window centered $L$ spatial units from the optimum.

Each of 200 assemblies (100 with $a = +0.9$, 100 with $a = -0.9$) were run, and analysis was carried out at the level of the assembly. The efficacy of a particular location for conservation interventions was determined by the percentage increase in the fraction of extant species in the conservation case ($\chi_{cons}$), compared to fraction of extant species in the the baseline (no intervention) case($\chi_{base}$): $\frac{(\chi_{cons}-\chi_{base})}{\chi_{base}} \times 100$.

Assemblages (16/200) where no species survived in the baseline case were excluded. Results are presented by fitting a GAM curve through the untransformed percentage increases for each of the skew directions. This identifies a clearly visible peak in both cases.

#### Demonstration of impact of changing $a$ parameter.

To demonstrate that the general lack of impact of changing the sign of the $a$ parameter, which we showed in the main text, is not due to a general insensitivity to $a$, we generated communities with $a$ values of -0.9, -0.7, -0.3, 0.3, 0.7 and 0.9, and then tested the critical rate of climate change as described above. As visible in the figure below, the results show a strong impact of $a$ on $v^*$ that is approximately symmetric about $a$ =0.

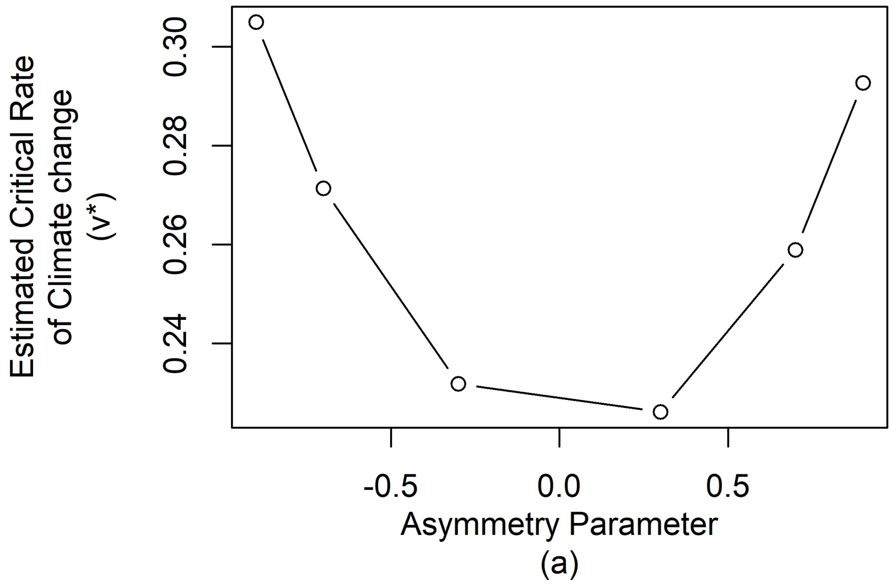


**Figure S3.2** Approximately symmetric impact of asymmetry parameter on the critical rate of climate change.
