## Supplementary material for "Schrödinger’s range-shifting cat: analytic predictions for the effect of asymmetric environmental performance on climate change responses": SI 4

### SI 4: Quantitative Performance of the Analytic Model

To further support our results, we examined how well the quantitative predictions of the analytic results corresponded to our simulation results. While many of the parameters in the simulation model align closely with the analytic model, the introduction of spatial heterogeneity, and the need to introduce density-dependent growth mean that there is not a perfect correspondence. To resolve this we use observations of our simulation model and the analytic model framework to identify reasonable approximations of the parameters used in our main results. We then test the predictions for the critical rate of climate change and the lag in our simulation models. While there is an inevitable degree of circularity in these analyses, in general they show a reasonable correspondence and provide support the use of our analytic model outside of the strict domain that it is defined within.

A R Markdown document that includes the code used to run the simulations and generate this document is available with the code supplement (`QuantitativeMatchTest.rmd`).

#### Estimating population growth rates, with and without climate change

To estimate the population growth rate without climate change $\lambda_0$, we first took an example assembled simulated community of 100 species as described in the other Appendices (using $a$ = -0.9, $\sigma = 0$). We then observed the increase in total population biomass of each species when the density dependent term was removed over 50 time steps. Dispersal between nodes carried on unaffected. This generated linear increases in log-biomasses (Figure S4.1a), through which we fit linear models to estimate the slope in each case. There was a moderate spread of estimated values across the 100 species, but the mean was approximately 4.96, which was the value we carried forward to later calculations.

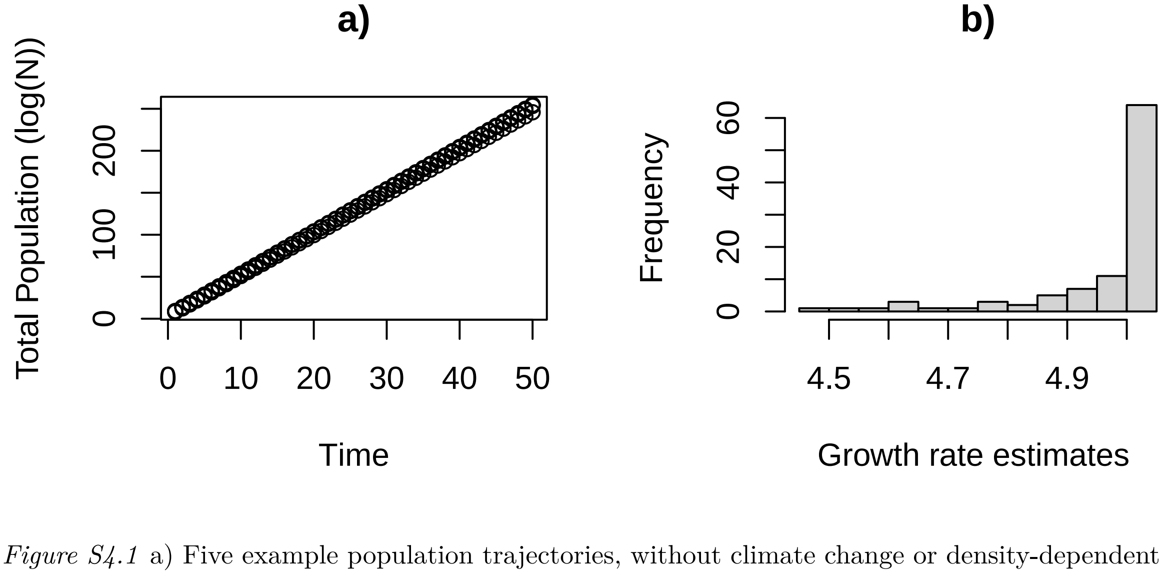


*Figure S4.1* a) Five example population trajectories, without climate change or density-dependent growth restraints, used to estimate $\lambda_0$. b) Histogram of estimated $\lambda_0$ values showing distribution of species responses.

In order to estimate $\lambda$, the growth rate with climate change, we repeated the estimation procedure, but with the introduction of climate change for 50 time steps. To allow for initial distribution adjustment, we fit our linear model through the final 25 time steps. We trialled this for a series of climate change rates $v$ up to 0.03, in which region climate change as a quadratic impact on population growth rate, as predicted by our model.

#### Inferring dispersal rates

To infer the effective dispersal rates in terms of the analytic model parameter $D$ in our spatial heterogeneous simulated communities, we use the expression derived in the main text for the reduction in growth rates with climate change, $\lambda = \lambda_0 - v^2/D$, rearrange to find $D = v^2/(\lambda_0 - \lambda)$ (Fig. S4.2b). We fit a linear model through the estimates of $D$ (excluding the two lowest values of $v$)to determine the value of $D$ when $v = 0$ (0.000775).

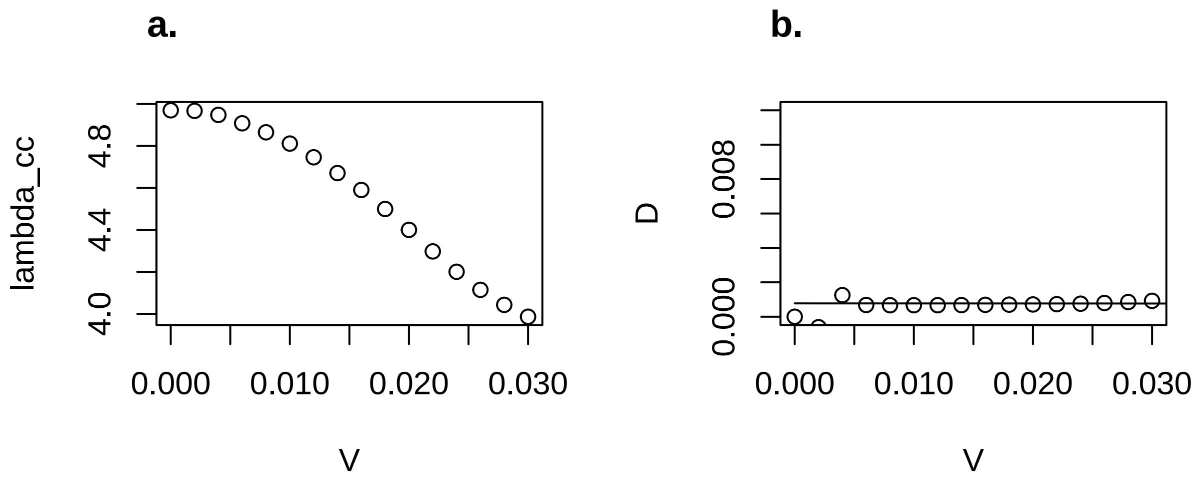


*Figure S4.2* a) Estimates of intrinsic growth rate in the presence of climate change $\lambda$, where climate change is introduced at various rates ($v$). b) inferred values of the dispersal rate $D$ using trials at different speeds of climate change.

#### Predicting $v*$

Then using the result from the main text $v^* = \sqrt{4D\lambda_0}$, and the estimate $D$ for the low $v$ case, we next estimate the expected critical value of climate change that leads to extinction as 0.124. This of the right magnitude but still substantionally lower than the 'true' value, which from simulations was found to be around $v = 0.35$.

#### Inferring $R_{max}$ and predicting lag.

Although a value of $R_{max}$ is specified in the simulation model (as 8), other changes in the model may not make this relation between models direct. We estimate the value of the maximum growth rate at the optimum $R_{max}$ using the expression

$$\lambda_0 = R_{max} - \frac{\sqrt{DR_{max}}}{w} + \frac{a^2 D}{4}$$

which we solve to find $R_{max}$ (assuming $w = 1$, to simplify):

$$R_{max} = \frac{1}{4}\left(2\sqrt{-aD^2 + D^2 + 4D\lambda_0} - aD + 2D + 4\lambda_0\right)$$

We then use these values to estimate the expected lag in time with climate change from:

$$\Delta^{time} \to \frac{w}{2\sqrt{DR_{max}}} + \frac{a^2 w \left(\sigma^2 \sqrt{DR_{max}} + 2Dw\right)}{4DR_{max}}$$

which since we do not here include weather ($\sigma = 0$), and maintain $w$ as 1, simplifies to:

$$\Delta^{time} \to \frac{1}{2\sqrt{DR_{max}}} + \frac{a^2}{R_{max}}$$

reaching:

$$\Delta^{time} = 8.17$$

Converted into lags in space, at $v = 0.1$, $\Delta^{space} = 0.817$, and at $v = 0.2$, $\Delta^{space} = 1.635$. The corresponding observed spatial lag values from the corresponding simulations were 0.478 and 1.01. The magnitude is approximately correct, but there is a significant discordance.
